# *Salmonella* cancer therapy metabolically disrupts tumours at the collateral cost of T cell immunity

**DOI:** 10.1101/2023.01.12.523780

**Authors:** Alastair Copland, Gillian M. Mackie, Lisa Scarfe, David A.J. Lecky, Nancy Gudgeon, Riahne McQuade, Masahiro Ono, Manja Barthel, Wolf-Dietrich Hardt, Hiroshi Ohno, Sarah Dimeloe, David Bending, Kendle M. Maslowski

## Abstract

Bacterial cancer therapy (BCT) is a promising therapeutic for solid tumours. *Salmonella enterica* Typhimurium (STm) is well-studied amongst bacterial vectors due to advantages in genetic modification and metabolic adaptation. A longstanding paradox is the redundancy of T cells for treatment efficacy; instead, STm BCT depends on innate phagocytes for tumour control. Here, we used distal T cell receptor (TCR) reporter mice (*Nr4a3-Tocky-Ifng-YFP*) and a colorectal cancer (CRC) model to interrogate T cell activity during BCT with attenuated STm. We found that colonic TILs exhibited a variety of activation defects, including IFN-γ production decoupled from TCR signalling, decreased polyfunctionality and reduced T_CM_ formation. Modelling of T-cell–tumour interactions with a tumour organoid platform revealed an intact TCR signalosome, but paralysed metabolic reprogramming due to inhibition of the master metabolic controller, c-Myc. Restoration of c-Myc by deletion of the bacterial asparaginase *ansB* reinvigorated T cell activation, but at the cost of decreased metabolic control of the tumour by STm. This work shows for the first time that T cells are metabolically defective during BCT, but also that this same phenomenon is inexorably tied to intrinsic tumour suppression by the bacterial vector.

## Introduction

Bacterial cancer therapy (BCT) is the antecessor of modern immunotherapy, having origins in the late 19^th^ Century, during which time U.S. physician William Coley used injections of *Streptococcus* to induce tumour regression in inoperable patients (Starnes, 1992). Yet it is only recently that there has been such a sharp resurgence of interest in BCT, owing to significant advances in genetic modification of bacterial vectors, coupled with safety and efficacy concerns regarding immune checkpoint blockade (ICB).

*Salmonella enterica* serovar Typhimurium (STm) is among the most-studied bacterial vectors for cancer therapy. As a metabolic ‘generalist’ (Dandekar et al., 2014), the bacterium can rapidly adapt to harsh tumour microenvironments, alongside potent tumour-homing abilities. We recently demonstrated in colorectal cancer (CRC) models that an attenuated STm mutant can reduce colonic tumour burden *in vivo* by imposing strong metabolic competition on the tumour, with particular reduction in amino acids, sugars and tricarboxylic acid (TCA) cycle intermediates, as well as specific targeting of tumour stem cells resulting in normalisation of tissue ontogeny (Mackie et al., 2021). The immunological consequences of these changes, however, are hitherto unexplored.

It is widely reported that STm induces a cascade of immune changes within the tumour microenvironment—such as widespread immune cell influx, pro-inflammatory cytokine production, dendritic cell (DC) maturation and natural killer (NK) cell activation—that is believed to support immune tumour-clearance (Yang et al., 2021). Indeed, monocyte depletion (Johnson et al., 2021), NK cell depletion (Lin et al., 2021), or *MyD88* deletion (Zheng et al., 2017) can wholly abrogate therapeutic success *in vivo*, underscoring the necessity of an innate-like response in this therapy.

Paradoxically, T cells appear to play little role in *Salmonella* BCT, despite their established role in tumour control. It has been reported for over two decades that tumour control is proportionally identical between wild-type (WT) mice and athymic nude/SCID mice after *Salmonella* treatment (Kaimala et al., 2014; Lin et al., 2021; Luo et al., 2001); indeed, many efficacy studies are conducted in athymic mice with xenograph tumours (Zhao et al., 2005; Zheng et al., 2017). Antibody depletion of CD4/CD8 T cells does not fully prevent *Salmonella*-mediated reduction in tumour volume (Lee et al., 2011), and adoptive transfer of anti-tumour, STm-generated T cells show no therapeutic capacity (Stark et al., 2009). Most strikingly, mice which have been cured of tumours by STm treatment do not possess an anti-tumour memory response upon re-challenge (Yoon et al., 2017). Taken together, T cells are likely redundant for *Salmonella* BCT.

In this study, we combined an autochthonous colitis-associated CRC mouse model using STm^Δ*aroA*^ (an attenuated aromatic amino acid auxotrophic mutant) with the *Nr4a3-Tocky-Ifng-YFP* kinetic TCR-cytokine reporter, which has recently been utilised to detect favourable responses to ICB (Elliot et al., 2021). This allowed unparalleled resolution of T cell activation dynamics during STm BCT. We found that T cells were indeed dysfunctional during STm BCT, showing profound decoupling of several activation nodes. This was not associated with any detectable defects in proximal T cell receptor (TCR) signalling, but instead a potent metabolic disruption and inability to sustain a conventional activation trajectory. T cell metabolic dysfunction was due solely to asparagine depletion by bacteria, leading to depleted c-Myc protein; reversal of this depletion restored T cell function. Critically, STm-mediated c-Myc suppression was also detected in the tumour itself, which dampened tumour stemness and survival, highlighting an important ‘double-edged sword’ for STm BCT in which tumour control by bacteria comes at the detriment of adaptive immunity. These findings provide a strong rationale for addressing a previously unknown cardinal defect in *Salmonella*-based cancer therapies to yield more successful clinical outcomes.

## Results

### *Salmonella* cancer therapy induces T cells with an aberrant activation signature

T cells are reported as redundant for *Salmonella*-based cancer therapies (Kaimala et al., 2014; Lin et al., 2021; Luo et al., 2001), and so we questioned whether they possessed canonical activation during BCT. To address this, we utilised an autochthonous colitis-associated colorectal cancer model (CAC) in *Nr4a3-Tocky-Ifng*-YFP (‘Tocky-GS’) (**Figure 1A**) mice. These mice contain a distal *Nr4a3* TCR reporter (Bending et al., 2018a, 2018b) consisting of a mutant mCherry protein (Subach et al., 2009) that transitions from Blue to Red form (t1/2 = 4hrs) upon TCR triggering, giving temporal insights into cell activation. This was used in conjunction with an *Ifng*-YFP reporter (Price et al., 2012), in order to probe T cell activity in tumours and lymphatic tissues (gating strategy shown in **Sup. Fig. 1A**). Following tumour induction, mice were orally gavaged with two doses (once per week) of an auxotrophic STm mutant (STm^Δ*aroA*^) which we and others have previously reported to reduce tumour burden in cancer models (Fensterle et al., 2008; Johnson et al., 2021; Mackie et al., 2021).

**Figure 1.**
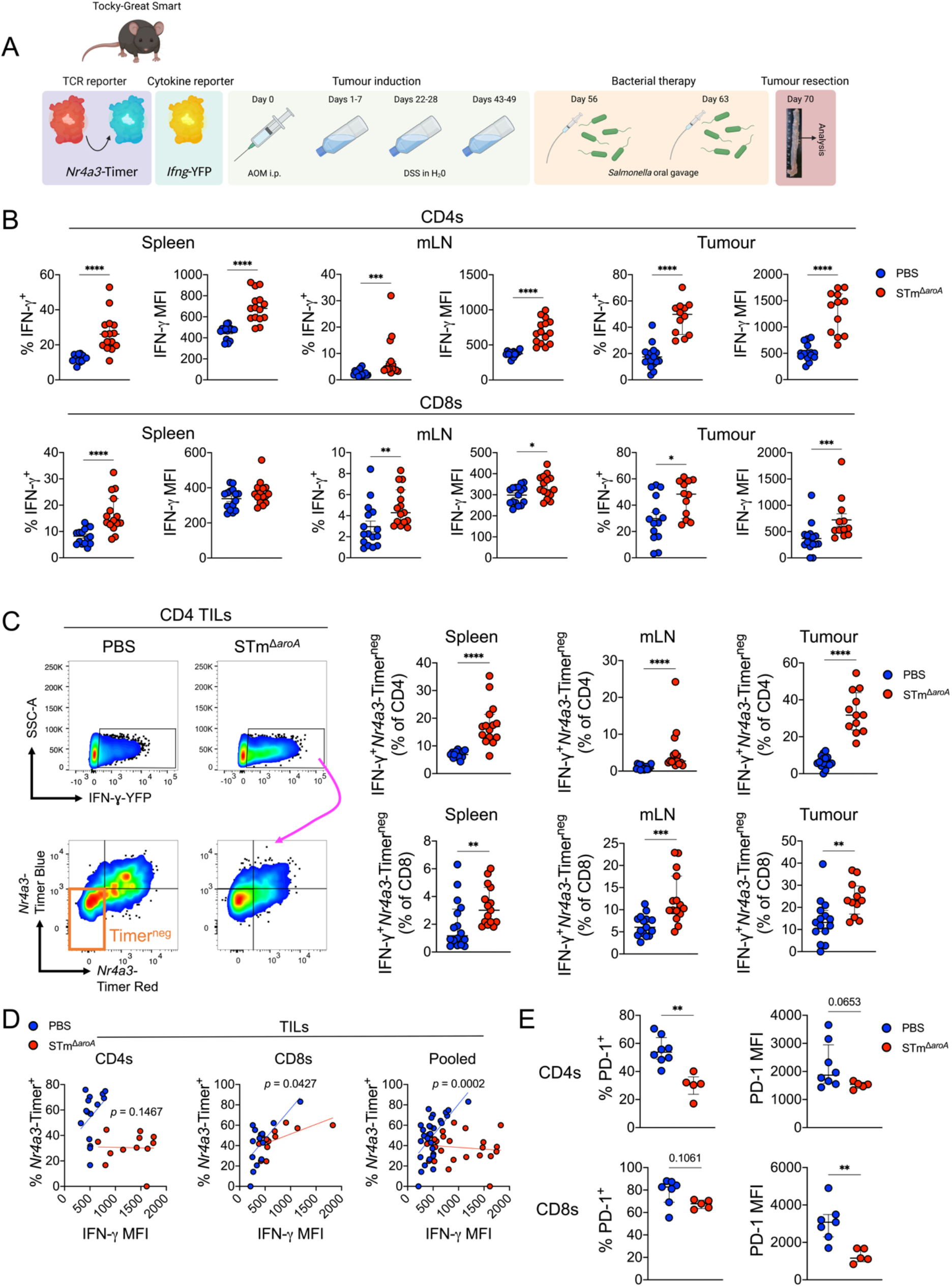
Attenuated *Salmonella* cancer therapy induces aberrant T cell activation that decouples IFN-γ from TCR signalling. **(A)** Schematic depicting the induction of colonic tumours in the CAC model using azoxymethane (AOM) and dextran sodium sulfate (DSS) and treatment with attenuated *Salmonella*. **(B)** Flow cytometric analysis of IFN-γ-YFP in CD4 and CD8 T cells, across spleen, mLN and tumour. Percentage positive and median fluorescence intensity (MFI) of the positive fraction are depicted. **(C)** *Left:* representative flow cytometry plot showing gating on IFN-γ^+^ T cells and sub-analysis of *Nr4a3*-Timer expression (Blue or Red form). *Right:* analysis of the IFN-γ^+^ and Timer^neg^ cells as a fraction of either CD4 or CD8 T cells. **(D)** Analysis of tumour IFN-γ^+^ T cells and a correlation between IFN-γ intensity (MFI) and frequency of TCR signalling (%*Nr4a3*-Timer^+^). Line depicts simple linear regression. **(E)** Flow cytometric analysis of PD-1 expression (percentage and MFI) within tumour IFN-γ^+^ T cells. Bars median average with interquartile range. Each point represents one mouse. Significance was tested by unpaired Mann-Whitney (*B, C, E*) or by linear regression of the two curves (*D*). Data are pooled from two similar experiments with similar data distribution.* = *p* < 0.05, ** = *p* < 0.01, *** = *p* < 0.001, **** = *p* < 0.0001

Flow cytometric analysis of mesenteric lymph nodes (mLN), spleen and tumour revealed that, as previously reported in other cancer models (Lee et al., 2008; Lin et al., 2021), *Salmonella* treatment induced high production of IFN-γ across all tissues in CD4 and CD8 T cells compared to the PBS control (**Figure 1B**). Strikingly, however, parsing of TCR reporter expression in reactive T cells showed that there was a strong bias towards a decoupling of IFN-γ expression from evidence of recent TCR signals, with STm generating significantly more IFN-γ^+^*Nr4a3*-Timer^neg^ T cells than the PBS control group (**Figure 1C**). Given that cytokine production and TCR signalling were being dissociated, we next tested whether the extent of IFN-γ expression (measured by MFI) correlated with the likelihood of TCR engagement (measured by *%Nr4a3*-Timer^+^) in TILs. As shown in **Figure 1D**, T cells in mice treated with PBS had a much stronger correlation between TCR engagement and magnitude of IFN-γ production, whereas *Salmonella* induced a pronounced decoupling of these aspects of T cell activation (*p* = 0.0002 PBS vs STm, pooled T cells).

To assess whether TCR engagement was indeed hampered during bacterial infection, PD-1 was used as a marker of TCR triggering. PD-1 is under strict control of the transcription factor NFAT (Martinez et al., 2015), which translocates to the nucleus upon TCR ligation. During T cell activation, PD-1 is usually co-expressed with *Nr4a3* (Elliot et al., 2021). As expected, PD-1 expression in both groups correlated tightly with evidence of TCR engagement (**Sup. Fig. 2**). Crucially, however, IFN-γ^+^ TILs in *Salmonella-treated* mice exhibited a reduction in expression of PD-1 when compared to the PBS control group, in keeping with the TCR reporter showing less evidence of TCR engagement (**Figure 1E**).

### T cells from infected tumours are hyporesponsive to conventional TCR-based activation

Given that T cells in STm-based BCT were being unconventionally activated, we hypothesised that T cells would be refractory to TCR stimulation. To this end, we employed a recently described *ex vivo* tumour platform (Voabil et al., 2021). Primary colonic tumour fragments were isolated and cultured in Matrigel to preserve the tumour architecture and microenvironment, thus permitting synchronous and polyclonal interrogation of activation dynamics. Tumours from PBS and *Salmonella-treated* mice were subjected to a-CD3 and a-CD28 stimulation *ex vivo* and *Nr4a3-Timer* was measured in T cells, with the hypothesis of dysfunctional activation (**Figure 2A**). CD4 T cells were analysed as they were the most abundant T cell type; CD8 T cells were scarce after culture in Matrigel (**Sup. Fig. 3A**).

**Figure 2.**
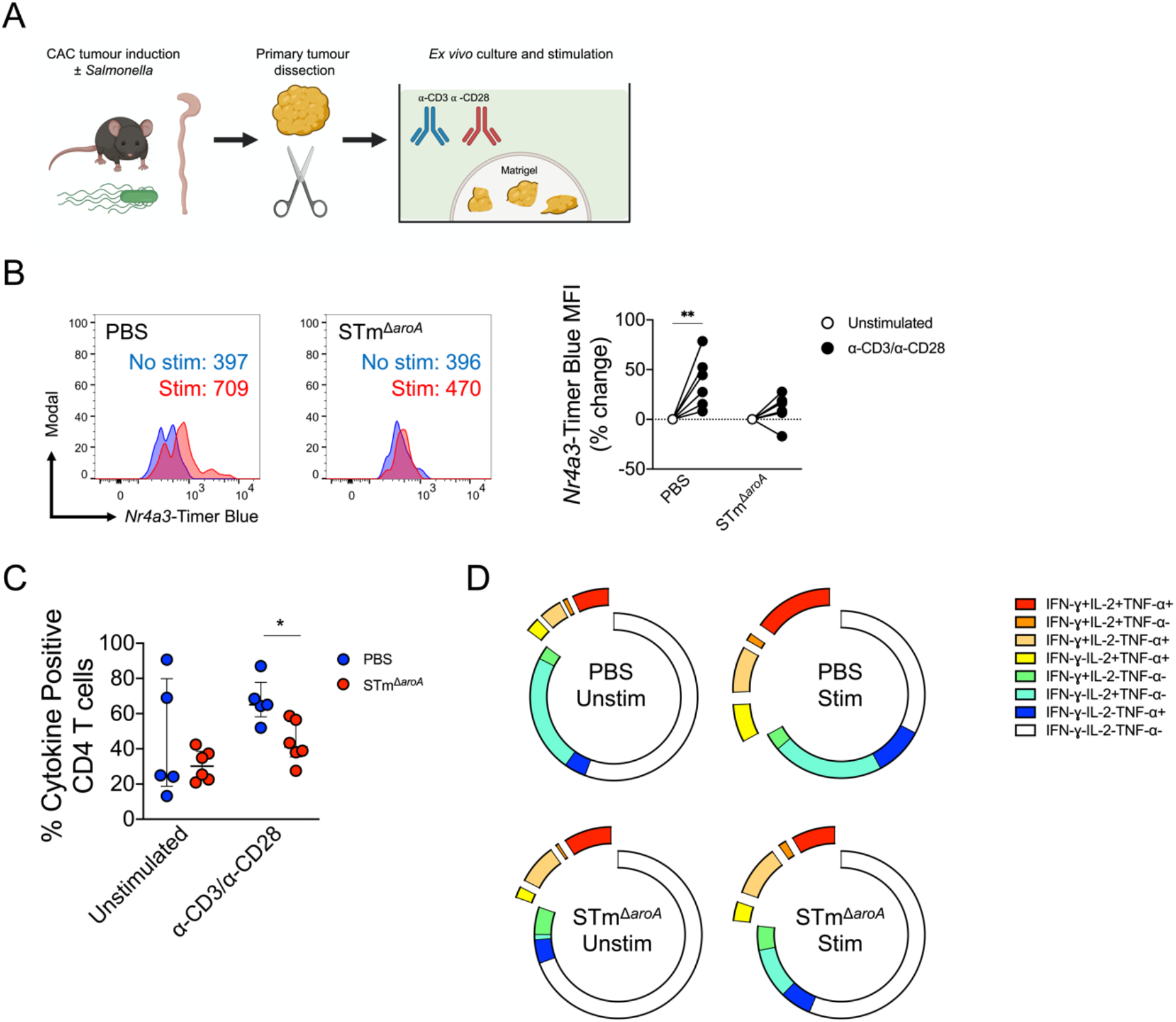
TILs from mice treated with *Salmonella* are functionally defective. **(A**) Schematic depicting dissection of primary tumours in the CAC model, followed by culture in Matrigel for *ex vivo* synchronous activation. **(B)** Primary TILs were stimulated with a-CD3/a-CD28 antibodies (1 μg/mL and 5 μg/mL, respectively) for 16h and then CD4 T cells were assessed for *Nr4a3*-Timer expression by flow cytometry, indicative of TCR-driven T cell activation. *Left:* Representative histograms showing T cell activation in PBS and STm-treated mice. *Right:* Pooled data from multiple mice showing % change in *Nr4a3*-Timer MFI upon activation. **(C)** Tumour fragments were activated with a-CD3/a-CD28 antibodies in the presence of brefeldin A (10 μg/mL) for 6h and concurrent cytokine production was measured by intracellular cytokine staining (ICS). Displayed is the total cytokine production in CD4 T cells. **(D)** Boolean analysis was performed on each possible cytokine combination in all four conditions. Shown are combined results from all mice depicting averages. Each point represents one mouse. Bars depict median average with interquartile range. Significance was tested by two-way ANOVA with Sidak’s post-test (unstimulated vs stimulated) (*B*) or by multiple Mann-Whitney tests with Holm-Sidak correction for multiple tests (C). *B* and *C* are from independent experiments. * = *p* < 0.05, ** = *p* < 0.01, *** = *p* < 0.001, **** = *p* < 0.0001

As can be seen in **Figure 2B**, polyclonal activation of CD4 T cells in primary tumours led to blunted *de novo Nr4a3-Timer* Blue expression, which is expressed immediately after triggering of the TCR complex. CD4 TILs from PBS-treated animals were able to up-regulate *Nr4a3*-Timer Blue (~40% increase, *p* < 0.01), whereas there was no significant change for the STm group. Next, we wished to know whether function was also affected. T cell cytokine polyfunctionality is a hallmark of optimal TCR engagement, and is beneficial against tumours and infectious diseases (Hart et al., 2018; Zhao et al., 2016). Tumour fragments were therefore activated and tested for concurrent Th1 cytokine production. As shown in **Figure 2C**, CD4 T cells from the STm-infected mice showed an overall blunted production of TCR-driven cytokine production, with fewer triple- and double-cytokine positive cells when analysed qualitatively (**Figure 2D**). These data confirmed that T cell activation was being actively blunted by *Salmonella* in the context of cancer therapy.

### T cell dysfunction is replicated by *Salmonella*-infected tumour organoids

We have previously observed strong concordance between the changes caused by STm *in vivo* and *in vitro*, using tumour organoids to explore changes in the metabolome and stem cell compartment during BCT (Mackie et al., 2021). Tumour organoids (small intestinal and colonic) were therefore generated and infected with STm for up to 48 hours, including a gentamicin wash-supplement step to kill bacteria outside of the intracellular tumour niche. This allowed for splenocytes and T cells to be directly co-cultured and assessed for activation within the tumour microenvironment (TME), or co-cultured with tumour-conditioned media (TCM) (**Figure 3A**). Since microbial TLR ligands can positively or negatively affect lymphocyte activation (Srinivasan and McSorley, 2007; Sturm et al., 2005), heat-killed (HK) STm was used as a control at an equivalent concentration to the live bacteria. T cell activation was assessed by the flow cytometric location and intensity of the *Nr4a3*-Timer (**Figure 3B**).

**Figure 3.**
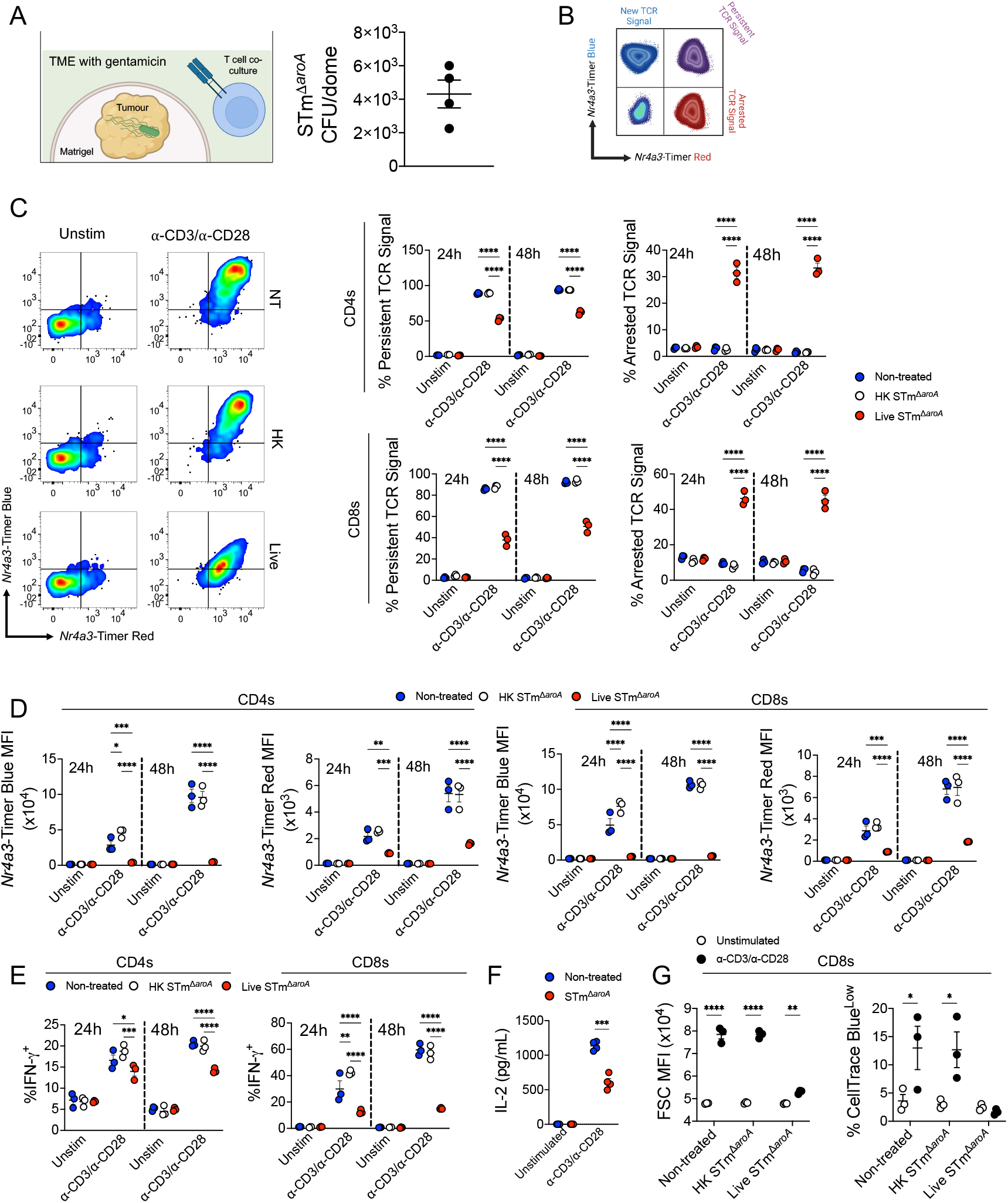
*Salmonella*-infected organoids recapitulate induction of T cell dysfunction and demonstrate arrested TCR activation. **(A)** *Left:* Schematic depicting the co-culture model of tumour organoid culture with T cells/splenocytes. Cells (1-2 x 10^6^ splenocytes) were either directly cultured with the tumour or with TCM. In direct cultures T cells infiltrate the Matrigel. *Right:* Representative intra-tumoural CFU counts after 48h culture, each dot represents one Matrigel dome in a 24-well plate. **(B)** Schematic of flow cytometric analysis of *Nr4a3-timer*. **(C)** Splenocytes were cultured directly with uninfected, HK-STm-treated or live STm-infected tumours for up to 48h ± a-CD3/a-CD28 antibodies (1 μg/mL and 5 μg/mL, respectively), and at each timepoint cells were aspirated for analysis by flow cytometry at the indicated time-points. *Left:* Representative plot depicting *Nr4a3-Timer* expression after stimulation in the three groups (Blue^+^Red^-^ = New TCR Signal, Blue^+^Red^+^ = Persistent TCR Signal, Blue^-^Red^+^ = Arrested TCR Signal). *Right:* Summary data showing locus of TCR reporter in CD4^+^ (top) or CD8^+^ (bottom) cells. **(D)** Summary analysis of *Nr4a3*-Timer Blue and Red MFI values. **(E)** Summary data for IFN-γ^+^ T cells at 24 or 48h. **(F)** Splenocytes were stimulated for 24h in TCM from NT or STm-infected tumours and IL-2 was measured in supernatants by ELISA. **(G)** Splenocytes were stained with CellTrace Blue and stimulated for 3 days in TCM from NT or uninfected tumours as described above. *Left:* Flow cytometric measurement of cell size (FSC-A) after stimulation. *Right:* Analysis of proliferating cells by gating on the CellTraceBluelow fraction (as a percentage of CD8^+^ cells). Bars depict means ± SEM. Each point represents one mouse or organoid infection. Statistical significance was tested by two-way ANOVA comparing conditions unstimulated or stimulated groups with Tukey’s post-test (*C, D, E, G*) or by two ANOVA with Sidak post-test comparing unstimulated to stimulated responses (*H*). Data are from *n* = 3 mice in three organoid infections and representative of more than five experiments (*C, D, E*), from *n* = 6 mice testing TCM from four infections (G), or *n* = 3 mice testing TCM from three independent experiments.* = *p* < 0.05, ** = *p* < 0.01, *** = *p* < 0.001, **** = *p* < 0.0001

As expected, T cells activated in the presence of non-infected or heat-killed STm-treated tumours were able to fully activate, as indicated by the high expression of both Timer Blue and Timer Red (>80% Timer Blue^pos^Timer Red^pos^, i.e. persistent TCR signalling), and only minor levels of Timer Blue^neg^Timer Red^pos^ (arrested TCR signalling) cells (**Figure 3C**). Strikingly, T cells activated in the presence of tumours infected with live *Salmonella* showed a pronounced failure to sustain activation (>40%, *p* < 0.0001 live STm vs non-treated and HK STm at 24 hours). This aborting of T cell activation was similarly reflected in the lower fluorescence intensity of Timer Blue and Timer Red, as measured by MFI (**Figure 3D**).

TCR signalling is intimately connected with cytokine production (Tao et al., 1997), and so we next assessed IFN-γ expression. As with TCR signalling, T cells activated in the milieu of uninfected or HK STm-treated tumours were able to produce high levels of IFN-γ (~50% IFN-γ^+^ CD8s at 48h), but there was a dampening of cytokine expression in lymphocytes exposed to STm-harbouring tumours (*p* < 0.0001 live STm vs NT and HK STm at 48h), particularly in CD8 T cells (**Figure 3E**). This was not limited to type 1 cytokines, as IL-2 secretion was similarly affected in T cells cultured with infected tumours (**Figure 3F**). Since IL-2 is a key cytokine for T cell proliferation and growth, we next analysed cell size and division after 3 days of culture. As shown in **Figure 3G**, CD8 T cells in the live STm group were smaller after stimulation, suggesting a lower proportion of cells undergoing mitosis. In accordance, these cells were unable to divide, as measured by dilution of tracer dye (*p* > 0.05, live STm unstimulated vs stimulated) (**Figure 3G**). Taken together, STm-infected tumours can disrupt multiple facets of T cell activation.

### RNAseq analysis shows *Salmonella*-infected tumours disrupt key genes involved in T cell metabolic reprogramming

We had established that T cells had hallmarks of defective activation both *in vivo* and *in vitro*. Therefore, we next sought to determine which signalling pathways were being disrupted by STm-infected tumours. Splenocytes were cultured with TCM (representing the TME) from infected or non-infected tumour organoids and T cells were polyclonally activated for 4 or 16 hours, representing early and late time-points for activation and *Nr4a3* expression. Following activation, CD4 T cells were FACS sorted to high purity and assessed for TCR signal, which showed that the arrested TCR signal was beginning to accumulate at this timepoint in the infected TCM condition (**Figure 4A**; 6.5% TCR-arrested in NT group, 28.1% TCR-arrested in STm group). CD4 T cells were then lysed and 3’ mRNA libraries were created and analysed by RNAseq Quant-Seq.

**Figure 4.**
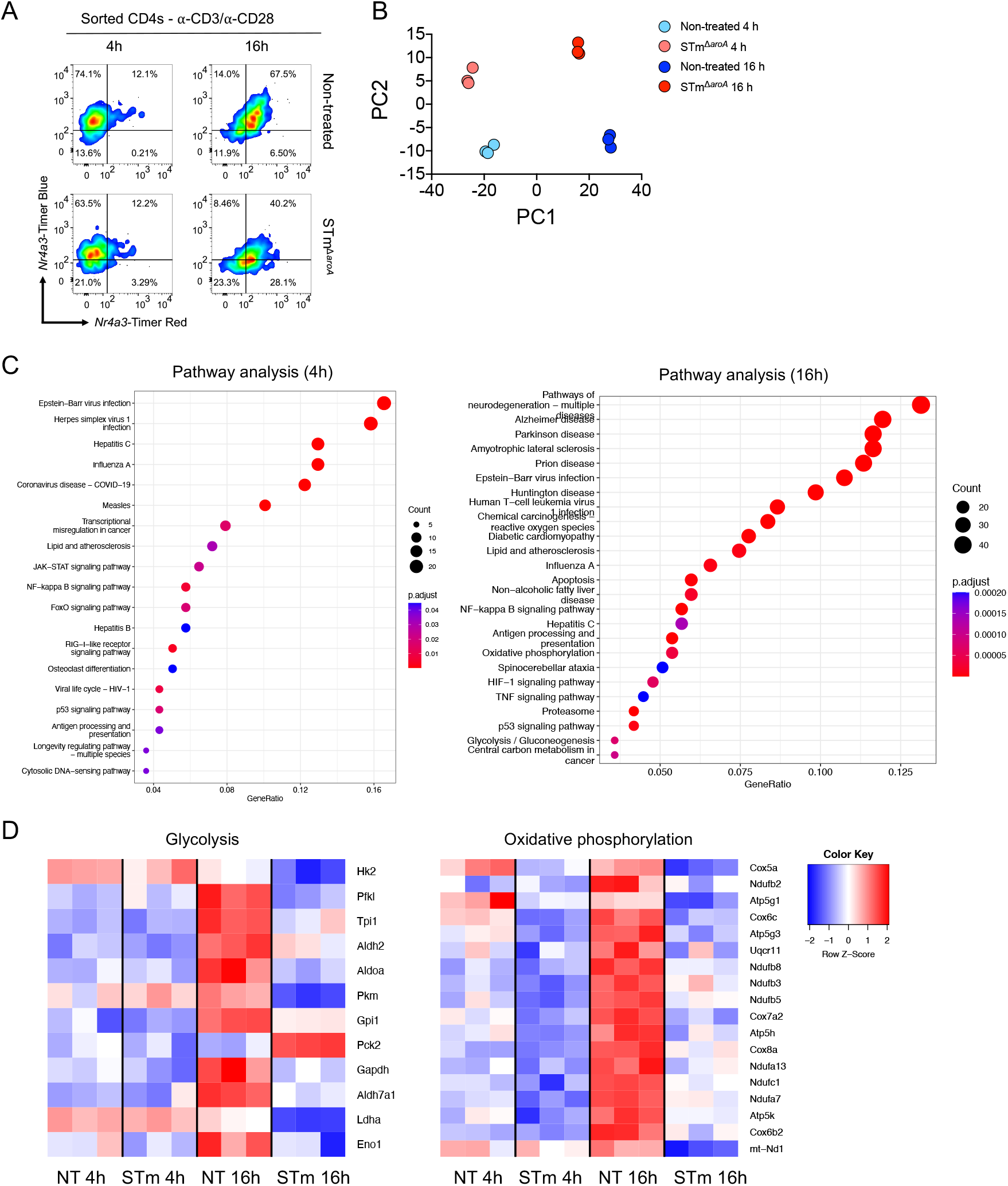
*Salmonella*-infected tumours down-regulate key metabolic genes in T cells, accompanied by an innate-like transcriptional signature. **(A**) Splenocytes (1 x 10^6^ per condition) were cultured with TCM from non-treated or STm-infected tumours for 4 or 16 hours, and then purified CD4 T cells were FACS sorted from these bulk splenocytes and *Nr4a3*-Timer expression was measured by flow cytometry. Data shown are a representative mouse. **(B)** 3’ mRNA libraries were generated from isolated CD4s and transcription was quantified using Quant-Seq. Shown is a Principal Component Analysis (PCA) displaying variance in the data set along PC1 and PC2. **(C)** DEGs were inputed into the KEGG database for pathway analysis for 4h (*left*) and 16h (*right*). **(D)** Heat-map of the DEGs involved in glycolysis (*left*) or oxidative phosphorylation (*right*) with Z-score key. Data are derived from *n =* 3 mice, representing one of two similar experiments. All DEGs were generated with a 1.5-fold change threshold.

Principal component analysis (PCA) showed strong separation of the four groups along PC1 and PC2 axes by both timepoint and tumour treatment (**Figure 4B**). Pathway analysis revealed a higher number of differentially expressed genes (DEGs) at the 16h timepoint compared to 4h timepoint, suggesting a progressive reprogramming of T cell phenotype (**Figure 4C**). Interestingly, the dominant pathways enriched in the STm group were involved in (i) innate antiviral responses (*Stat1, Stat2, Irf7, Irf9*) and (ii) metabolic reprogramming, namely glycolysis and oxidative phosphorylation (OXPHOS). We reasoned that TLR activation of T cells was likely responsible for the innate-like signature being enriched by exposure to STm-infected tumours, and therefore we turned our attention to the metabolic pathways. Subanalysis of metabolism-related DEGs revealed a striking suppression of multiple genes involved in glycolysis (*Hk2, Pfkl, Pkm, Ldha, Eno1*) by *Salmonella*-infected tumours at 16h; this was paralleled by dampening of gene expression in the OXPHOS pathway (*Cox5a, Atp5g1, mt-Nd1*) at the same time-point (**Figure 4D**). T cells exposed to non-infected tumours demonstrated normal metabolic reprogramming during this late-stage activation, with substantial upregulation of glycolysis and OXPHOS genes.

### T cells exposed to *Salmonella*-infected tumours have impaired metabolic capacity

Since RNA transcriptome analysis showed several enzymes and complexes in glycolysis and OXPHOS were suppressed by STm, we next focused on the metabolic features of T cell activation when cells were exposed to infected tumours.

First, it was tested whether inhibition of glycolysis could recapitulate the antagonism of TCR signalling by STm-treated tumours, i.e. *Nr4a3-Timer* arrest (Timer Red accumulation). To this end, T cells were treated with TCM from uninfected (nontreated; NT) or STm-treated tumours containing up to 20 mM 2-deoxy-D-glucose (2-DG), a potent inhibitor of glycolysis which leads to accumulation of 2-deoxy-d-glucose-6-phosphate (2-DG-6P). As predicted, addition of 2-DG to TCM was able to potently arrest T cell activation, as evidenced by a dose-dependent increase in arrested TCR signals in CD4s and CD8s when cultured in NT TCM (**Figure 5A**). At the highest dose of 2-DG, T cells in the NT tumour group were virtually identical to the STm tumour group in terms of arresting of the TCR reporter. The addition of 2-DG to the TCM from infected tumours was able to enhance the arrested TCR signalling even further (*p* < 0.01, CD4 and CD8 stimulated: vehicle vs 20 mM 2-DG). Similar results were found for IFN-γ expression: in CD8s, 2-DG suppressed cytokine production in the NT tumour group to levels seen in the STm tumour group (**Figure 5B**).

**Figure 5.**
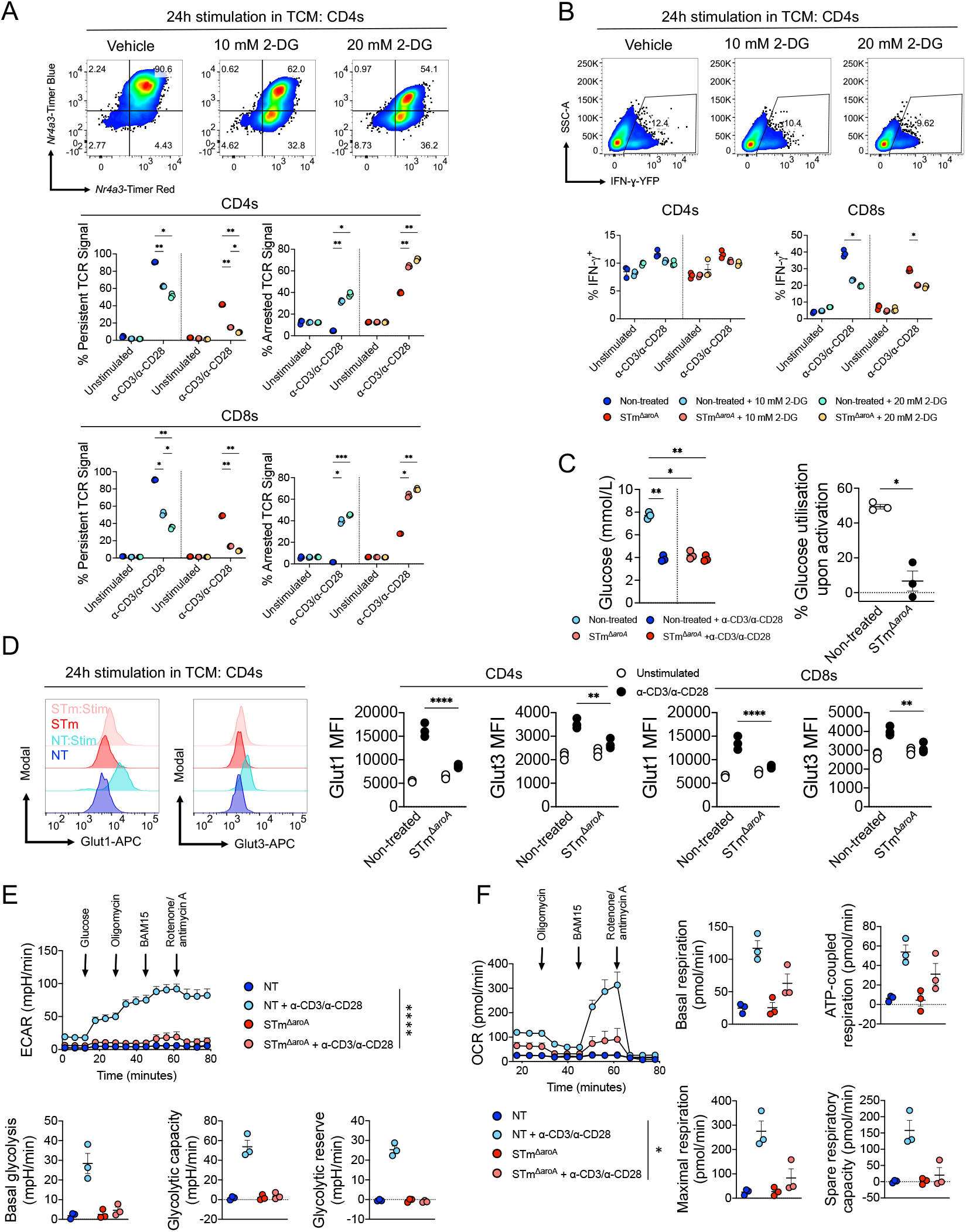
T cell metabolic paralysis is induced by *Salmonella*-infected tumours. **(A & B)** Splenocytes (1 x 10^6^/well) were cultured with TCM from non-treated or STm-infected tumours, alongside 10 - 20 mM 2-DG, or vehicle control (water) added at the beginning of the experiment. Indicated groups were then activated with a-CD3/a-CD28 antibodies (1 μg/mL and 5 μg/mL, respectively) and assessed for *Nr4a3*-Timer loci **(A)** and IFN-γ expression **(B)** in CD4^+^ and CD8^+^ T cells. Blue^+^Red^+^ = Persistent TCR Signal, Blue^-^Red^+^ = Arrested TCR Signal. **(C)** Splenocytes were cultured and activated for 24h in TCM from NT or STm-treated tumours. Supernatants were then tested (*left*) for glucose concentration (Sinocare), and percent utilisation was calculated (*right*). **(D)** Splenocytes were cultured and activated for 24h in TCM from NT or STm-treated tumours. After 24h, cells were fixed and permeabilised. Cells were stained with either rabbit anti-Glut1 or anti-Glut3, followed by anti-rabbit-APC to reveal total Glut1/3 levels. *Left*: Representative histogram showing flow cytometric levels of Glut1 and Glut3 gated on CD4 T cells. *Right:* Compiled results from *n* = 3 mice. **(E-F)** Splenocytes were cultured in TCM from NT or STm-treated tumours for 48h and then analysed on Agilent Seahorse XF analyser for ECAR **(E)** and OCR **(F).** Data are representative of two (A, B) or one (C) independent experiments testing *n* = 3 mice splenocytes on TCM from two infections, or one experiment testing TCM from two independent infections on *n* = 3 mice (D, E, F). Statistical significance was tested by two-way ANOVA comparing all groups with Sidak’s post-test, displaying the stimulated in-group comparisons (A, B), or one-way ANOVA with Tukey’s post-test and (C), two-way ANOVA with Sidak’s post-test between unstim vs stim (D, E) and one-way ANOVA with Tukey’s post-test, displaying the stimulated group only. * = *p* < 0.05, ** = *p* < 0.01, *** = *p* < 0.001, **** = *p* < 0.0001

Next, splenocytes were cultured in TCM from NT or STm-infected tumours and T cell activation was induced for 24h, followed by quantification of glucose utilisation in the culture (**Figure 5C**). In cells cultured with TCM from NT tumours, there was an approximate 50% reduction in glucose (*p* < 0.01) upon T cell activation. In cells cultured with TCM from STm-infected tumours, however, there was no detectable glucose utilisation, consistent with the previous observation of impaired glycolytic enzyme expression. STm treatment of organoids itself depletes glucose (Mackie et al., 2021), however the levels remaining (approximately 4mM) should be sufficient for activation, and supplementation of excess glucose could not restore the phenotype (**Sup. Fig. 4A**). Since Glut1 and Glut3 are the major glucose transporters in T cells and are rapidly upregulated following activation (Macintyre et al., 2014), we assessed their expression following activation in TCM from infected or uninfected tumours. As shown in **Figure 5D**, there was a severe blunting of the up-regulation of Glut1/Glut3 in response to STm, whereas T cells cultured in normal tumour media were able to strongly upregulate these glucose transporters.

To functionally explore these observations, we next performed analysis of glycolysis and OXPHOS by measuring the extracellular acidification rate (ECAR) and oxygen consumption rate (OCR), respectively by extracellular flux analysis. As shown in **Figure 5E**, T cells activated in media from STm-infected tumours demonstrated a severely blunted increase in ECAR when glucose was provided (*p* < 0.0001 NT stim vs STm stim), indicating diminished glycolytic capacity. Furthermore, following injection with electron transport chain (ETC) inhibitors, a strong trend for impaired overall glycolytic capacity and glycolytic reserve were also observed. Similar results were found for OXPHOS: the STm group demonstrated decreased maximal OCR upon injection with the mitochondrial uncoupler BAM15, when compared to the NT group (*p* < 0.05 NT stim vs STm stim). Strong trends were also observed for STm-mediated decreases in basal respiration, ATP-coupled respiration and spare respiratory capacity. Together with the RNAseq analysis, these results show STm induces significant impairment of T cell metabolic reprogramming.

### Activated T cells exposed to *Salmonella*-infected tumours have an intact TCR signalosome but impaired levels of c-Myc

The relationship between TCR signalling and T cell metabolic reprogramming is bi-directional: (i) “top-down”, in that TCR signalling drives metabolic reprogramming to meet the energetic demands of a fully activated lymphocyte, granting optimal effector functions, and (ii) “bottom-up”, describing how changes in metabolic effectors can modulate TCR signalling and the activation trajectory of the cell (Shyer et al., 2020). Using this framework, we asked the question: are STm-infected tumours preventing T cell metabolic reprogramming due to inhibition of TCR signalling (“top-down”), or are they affecting a metabolic effector directly, so that the T cell is unable to sustain TCR-driven activation (“bottom-up”)?

First, we assessed at one of the earliest phosphorylation events upon TCR triggering: Lck phosphorylation (Dutta et al., 2017). Splenocytes were cultured directly with infected or non-infected tumours for 24h and levels of Lck phosphorylation were detected by flow cytometry after 15 mins a-CD3/a-CD28 stimulation. Surprisingly, T cells cultured with infected tumours were able to activate this kinase to a similar extent as controls (**Figure 6A**). Downstream of proximal phosphorylation events at the TCR complex is the flux of calcium that leads to NFAT translocation into the nucleus. This was measured by culturing splenocytes with tumours and then briefly pulsing with the ionophore, ionomycin. As seen in **Figure 6B**, T cells cultured in all conditions were fully competent at inducing calcium flux, as determined by ratiometric measurement of two cytosolic calcium dyes (*p* = non-significant, ionomycin treatment NT vs STm in CD4s).

**Figure 6.**
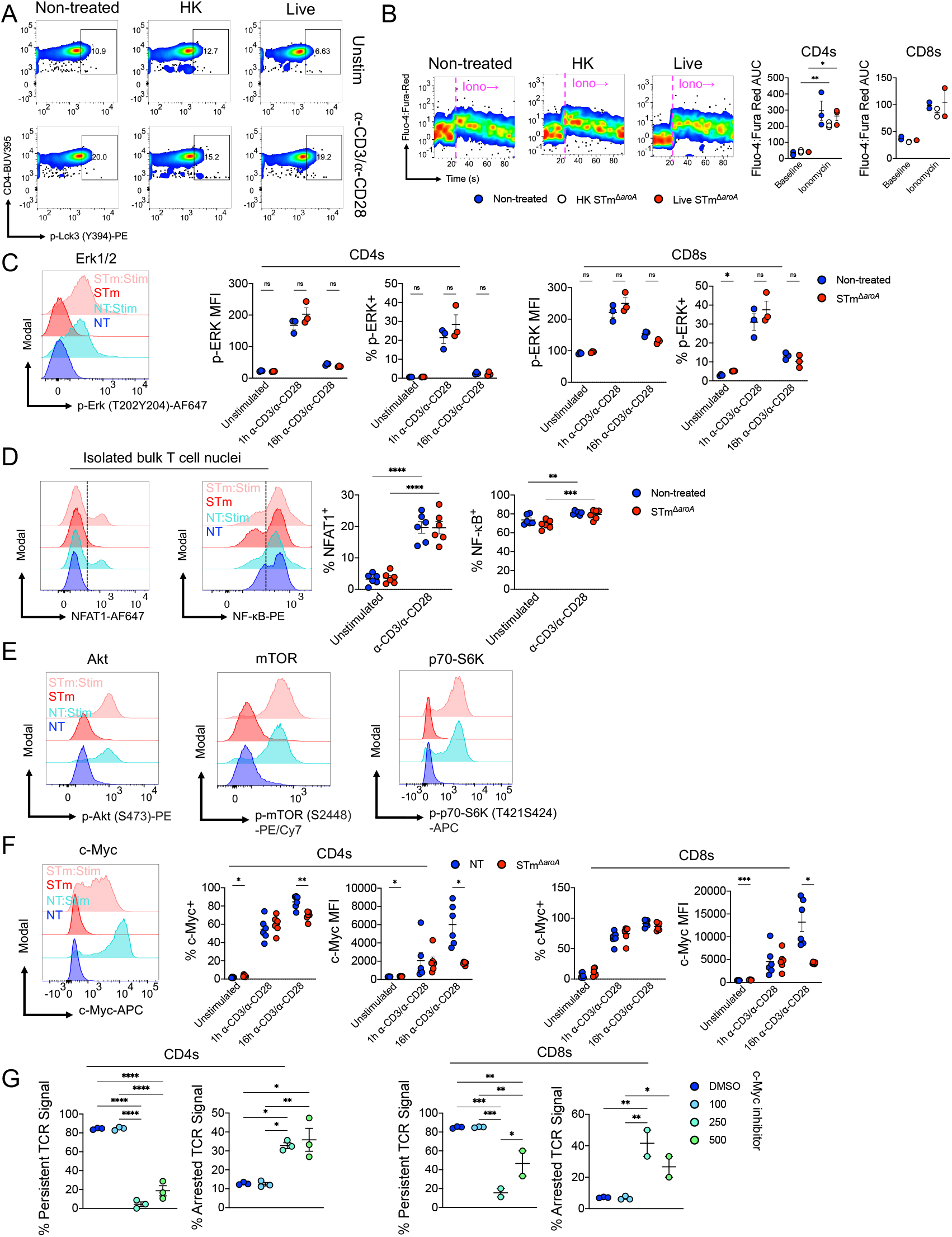
c-Myc is selectively inhibited by STm-treated tumours but the TCR signaolsome remains intact. **(A)** Splenocytes were cultured directly with non-treated, HK STm-treated or live STm-infected tumours for 24h. Cells were then stimulated for 15 mins with a-CD3/a-CD28 antibodies (1 μg/mL and 5 μg/mL, respectively) and then stained, fixed and permeabilised according to the PhosFlow protocol (see Methods). Cells were stained for Lck phosphorylation (Y394) by flow cytometry. Data shown are gated on CD4^+^ T cells. Gate was determined by a staining control. **(B)** Splenocytes were cultured directly with NT, HK STm-treated or live STm-infected tumours for 24h. Cells were then stained with lineage markers and Fluo-4/Fura-Red (calcium dyes), and run on a flow cytometer to measure baseline calcium. Ionomycin (1 μg /mL) was added after ~25 seconds, and calcium flux was measured by a ratio of Fluo-4:Fura-red and calculating the area under curve (AUC) before and after treatment. *Left:* Representative plots showing calcium flux for each group, gated on CD4 T cells. *Right*: Pooled data from *n* = 3 (CD4) or *n* = 2 (CD8) mice. **(C)** Splenocytes were activated in TCM for 1 or 16h with a-CD3/a-CD28. Cells were stained according to the PhosFlow protocol, and p-Erk (T202/Y204) was measured. *Left:* representative histogram showing induction in both activated groups. *Right*: Pooled data from *n* = 3 mice. **(D)** Purified bulk T cells (>98% purity) were cultured in NT or STm TCM for 24 hours. 1 hour before the end of the culture, a-CD3/a-CD28 was added to the indicated groups to measure nuclear translocation. Nuclei were then separated and analysed by flow cytometry for NFAT-1 and NF-κB. *Left:* Representative histograms showing NFAT1 and NF-κB translocation after activation. *Right:* Pooled data from 3 experiments, *n* = 6 mice. **(E)** Representative flow cytometry histograms of p-Akt (S473), p-mTOR (S2448) and p-p70-S6K (T421/S424). Cells were stimulated in NT and STm TCM for 1h and processed as previously described. Data representative of *n* = 3 mice. **(F)** Splenocytes were cultured in TCM for 1 or 16h with a-CD3/a-CD28. Cells were stained, fixed, permeabilised and stained with rabbit anti-c-Myc, followed by anti-rabbit APC. *Left*: Representative histogram after 1h stimulation. *Right*: Data pooled from two experiments, *n* = 6 mice. **(G)** Splenocytes were cultured in TCM from NT or STm-infected tumours and activated with a-CD3/a-CD28. At the beginning of culture, some conditions were treated with c-Myc inhibitor 10058-F4 or equivalent amount of DMSO. *Nr4a3-Timer* locus was measured by flow cytometry. Data shown are pooled from two (*B, F*), three (*D*) or one (*E, G*) independent experiments, or representative of two/three independent experiments (*A, C*, respectively). Bars depict means ± SEM. Statistical significance was tested by twoway ANOVA with Sidak post-test between treatments (*B, D*), time-points (NT vs STm) (*C, F*), or one-way ANOVA with Tukey’s post-test (G). * = *p* < 0.05, ** = *p* < 0.01, *** = *p* < 0.001, **** = *p* < 0.0001

While calcium flux drives NFAT-dependent T cell activation, mitogen-activated protein kinases (MAPKs) induce the second axis: the AP-1/NF-κB-dependent signalling branch (Rincon and Flavell, 1996). Erk phosphorylation was therefore measured in an activation time-course from 1-16h. As shown in **Figure 6C**, T cells exposed to TCM from both infected and non-infected tumours could induce robust Erk phosphorylation at 1h, which declined by 16h. With calcium flux and Erk phosphorylation intact, we next interrogated the immediate downstream transcription factors: NF-κB and NFAT. Purified bulk T cells were cultured in TCM from NT and STm-treated tumours, and then activated for 1h before staining for nuclear NFAT1 and NF-κB, according to a previously established protocol (Gallagher et al., 2021, 2018). In keeping with the upstream effectors, both NFAT1 and NF-κB were able to undergo equivalent nuclear translocation in both conditions, with approximately 20% T cells being NFAT1 ^+^ and 80% being NF-κB^+^ irrespective of tumour infection status (**Figure 6D**).

Aside from the NF-κB and NFAT1 axes, signalling via Pi3K-Akt-mTOR is also critical for full metabolic reprogramming and immune functionality (Pollizzi and Powell, 2015). Hence, activation status of this signalling pathway was measured. Surprisingly, three major proteins in this axis were similarly activated in T cells cultured with TCM from uninfected or infected tumours (**Figure 6E** & **Sup. Fig. 5**). In fact, there was a significant increase in mTOR activation at 16h in the STm group compared to the NT group in both CD4s and CD8s (NT: ~65% p-mTOR^+^, STm: ~80% p-mTOR^+^ in CD4s; **Sup. Fig. 5B**).

Finally, we investigated intracellular levels of c-Myc protein, since c-Myc is a master controller of cellular metabolism (Wang et al., 2011). Unexpectedly, while T cells activated in TCM from infected tumours could up-regulate c-Myc to a certain extent, this was substantially dampened compared to the NT control (*p* < 0.05 NT vs STm in CD4s and CD8s) at late-stage activation (**Figure 6F**), coinciding with the observation of impaired expression of glycolytic enzymes at the same time-point. To test whether c-Myc inhibition could replicate the termination of TCR signalling and T cell activation, T cells were activated after treatment with a small molecule c-Myc inhibitor, and *Nr4a3*-Timer was measured. As **Figure 6G** shows, c-Myc inhibition was able to significantly increase the percentage of arrested TCR signals in CD4s and CD8s, with a reciprocal drop in those with a persistent TCR signal.

Since c-Myc is a regulator of T cell memory formation (Haque et al., 2016; Nozais et al., 2021), we also looked *in vivo* at the memory phenotype of IFN-γ^+^ T cells in lymphatic tissue. Central memory T cells (T_CM_) are much more potent than effector memory T cells (T_EM_) in tumour control, and this ratio is an important biomarker for cancer survival (Liu et al., 2020). We detected an increase in TEM-like CD4 T cells in the spleen and mLN, and a corresponding decrease in T_CM_-like cells in STm-treated mice (**Sup. Fig. 5A** & **5B**, respectively), in keeping with T cell dysfunction. Taken together, these data revealed a “bottom-up” defect in T cell activation induced by STm: an intact TCR signalosome, but defective c-Myc expression leading to an inability to sustain canonical T cell activation.

### Reversal of tumoural asparagine depletion by *Salmonella* restores T cell function but loses direct control of cancer stem cells

Thus far, we had established that STm-infected tumours were suppressing T cell function by highly selective targeting of the metabolic controller, c-Myc. The next question we asked was, what is causing c-Myc suppression? C-Myc is a highly desirable drug target for tumours, given its role in promoting cancer stem cells, cell growth and proliferation. Clinical use of *E. coli-derived* L-asparaginase to deplete asparagine has a suppressive effect on c-Myc (Soncini et al., 2020), and like *E. coli, Salmonella* also encode asparaginases (Torres et al., 2016). To this end, we generated an attenuated STm mutant lacking *ansB*, which encodes a type II L-asparaginase, and used this strain to infect tumours both *in vivo* and *in vitro* (**Figure 7A** and **Figure 7B**).

**Figure 7.**
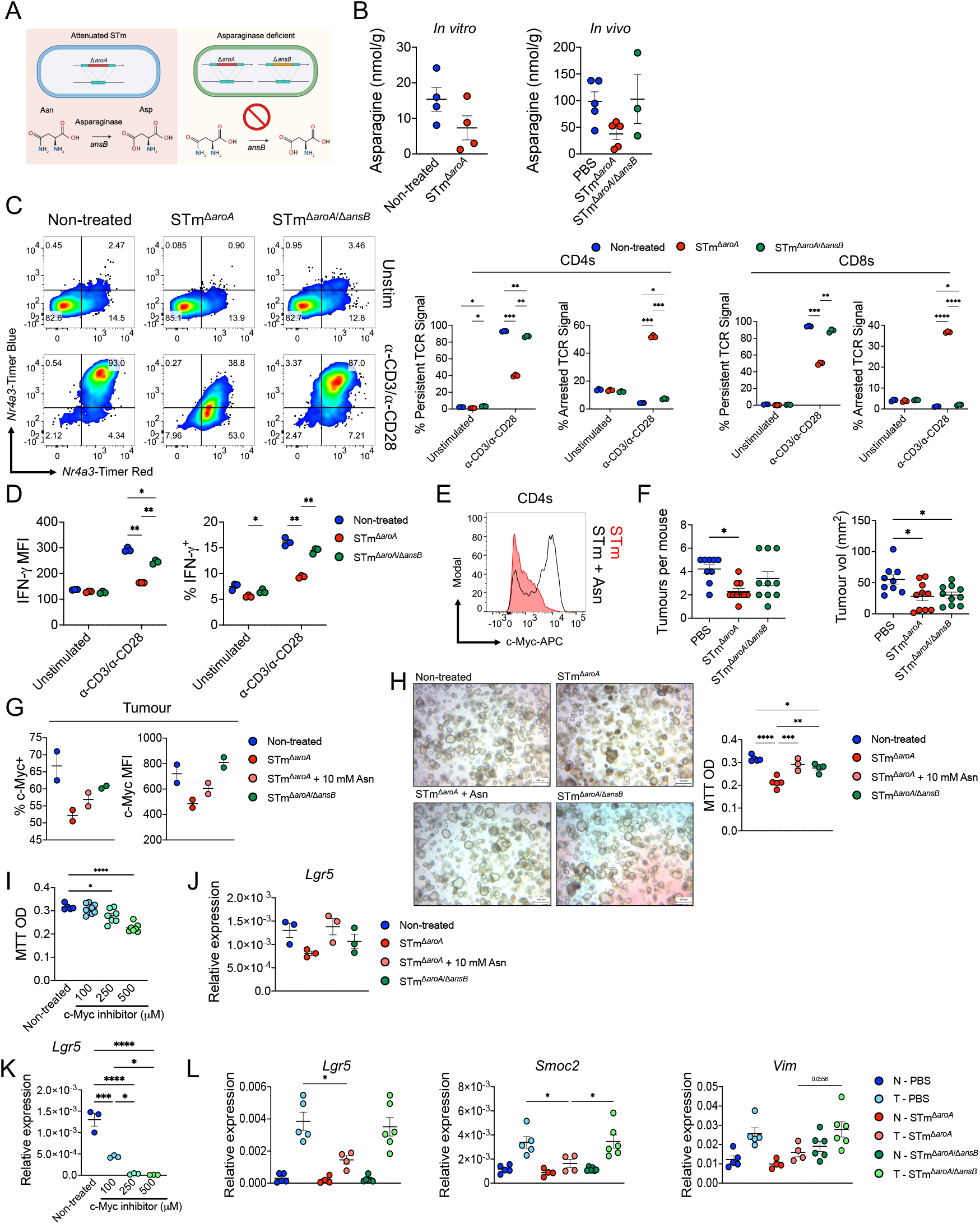
Deletion of bacterial asparaginase *ansB* restores T cell function to the loss of tumour stem cell control. **(A)** Schematic depicting the asparaginase-deficient attenuated STm mutant and the impaired catalytic reaction. **(B)** Asparagine was detected by GC-MS analysis of metabolites in tumour organoids (*left*) after 24h infection and after oral 6 doses of STm therapy *in vivo (right*) (Dataset taken from Mackie et al., 2021). **(C)** Splenocytes were cultured with TCM from tumours infected with aroA or ansB and activated with a-CD3/α-CD28 for 24h. *Left:* representative flow cytometric plot showing *Nr4a3*-Timer, gated on CD4 T cells. *Right:* pooled data from multiple mice, showing the Timer loci in relation to TCR signalling in CD4 and CD8 T cells. TCM pooled from two independent infections, *n* = 3 mice. **(D)** IFN-γ expression in the same experiment, showing pooled responses gated on CD4 T cells. **(E)** c-Myc levels in CD4 T cells after supplementation with 0.5 mM asparagine, as measured by intracellular flow cytometry. Representative of one of *n* = 2 mice. **(F)** CAC was induced in C57BL/6 mice using AOM/DSS followed by weekly oral gavage with 5 x 10^9^ CFU of STm^Δ*aroA*^, STm^Δ*aroA*/Δ*ansB*^ or PBS control for 6 weeks. Number of tumours per colon (tumour burden) and tumour volume were assessed. Each dot represents one mouse (*n* = 10 per STm group, 9 PBS group). (Data of NT v’s STm^Δ*aroA*^ taken from Mackie et al., 2021). **(G)** Tumour organoids were infected with STm^Δ*aro*A^ STm^Δ*aroA*^ + 10 mM Asn or STm^Δ*aroA*/Δ*ansB*^ for 24h then stained for intracellular c-Myc. Each dot represents one tumour line infection. **(H-I)** Tumour organoids were either infected with the indicated STm mutant with or without asparagine supplementation or with a c-Myc inhibitor for 24 hours. Organoid viability/metabolic capacity was assessed by MTT assay. Each dot represents an independent well. Representative of *n* = 2 independent experiments using two different *Apc^min/+^*-derived tumour organoid lines. **(J-K)** RNA was isolated from organoids in H-I and qPCR for *Lgr5* performed. **(L)** RNA was isolated from normal colon (N) or tumour (T) from experiment in (*F*) and qPCR analysis of indicated stem/mesenchymal transcripts performed. Each dot represents tumour from one mouse, data representative of >3 experiments for the PBS and STm^Δ*aroA*^ groups and 1 experiment for STm^Δ*aroA*/Δ*ansB*^ group. (Data of NT v’s STm^Δ*aroA*^ taken from Mackie et al., 2021). Bars depict means ± SEM or median with interquartile range. Statistical significance was tested by two-way ANOVA with Sidak’s post-test (*C, D*), one-way ANOVA with Tukey’s or Dunnett’s post-test (*F, H, I, J, K*), or Kruskal-Wallis with Dunn’s post-test (L). * = *p* < 0.05, ** = *p* < 0.01, *** = *p* < 0.001, **** = *p* < 0.0001

Remarkably, infection of tumour organoids with STm^Δaro*A*/Δ*ansB*^ was able to completely reverse the arrest of TCR signalling when T cells were activated in their TME, with a near-complete reversion of persistent TCR signalling to comparable levels as those in the NT group (**Figure 7C**). The bolstered TCR signalling was also reflected in IFN-γ production by these cells, with CD4 T cells gaining a significant boost compared to the STm^Δaro*A*^ control (*p* < 0.01 percentage and MFI) (**Figure 7D**). To confirm that this restorative effect was due to asparagine, TCM from STm^Δaro*A*^-infected tumours was spiked with asparagine at various concentrations. Consistent with the asparaginase-deficient mutant, asparagine supplementation was able to reverse the suppression of TCR signalling and IFN-γ production by STm (**Sup. Fig. 7**), alongside restoration of c-Myc expression (**Figure 7E**) to normal levels. Aspartic acid or glutamine were unable to restore T cell activation (**Sup. Fig. 7**).

With T cell function restored, we wanted to know how this would affect tumour load *in vivo* using the CAC model. Mice were treated with either PBS, STm^Δaro*A*^ or STm^Δaro*A*/Δ*ansB*^ for 6 weeks and then assessed for colonic tumour loads. As previously shown (Mackie et al., 2021), STm^Δaro*A*^ reduced the number and size of tumours in our model, but surprisingly the STm^Δ*aroA*/Δ*ansB*^ strain was unable to improve upon this outcome (**Figure 7F**). Questioning why this might be, we hypothesised that the same mechanism suppressing T cell function (c-Myc depletion) was important for direct control of the tumour by *Salmonella*, and could explain our previous findings of the bacteria imposing direct metabolic and stem-modulating effects on the tumour (Mackie et al., 2021). To explore this, we first measured c-Myc levels in the tumour itself and found that c-Myc was indeed lowered by STm^Δ*aroA*^, which could be reverted by exogenous asparagine or the asparaginase-deficient mutant (**Figure 7G**). Using an MTT assay to quantify the metabolic activity of the tumours, we found that STm^Δ*aroA*^ infection significantly reduced the size and metabolic activity of tumour organoids (*p* < 0.001 NT vs STm^Δ*aroA*^), which was reversed again by addition of exogenous asparagine or the asparaginase-deficient mutant (**Figure 7H**). To link this to c-Myc, we also then tested a small molecule inhibitor to see if this could replicate the effects of STm^Δ*aroA*^ infection, which was found to be the case (**Figure 7I**). We next assessed the effect of excess asparagine or asparaginase-deficient STm on tumour organoid expression of the stem cell marker *Lgr5*, as we have previously reported that STm^Δaro*A*^ causes a reduction of multiple indications of tumour stemness, including expression of *Lgr5*. There was a strong trend for a reduction in *Lgr5* expression by STm^Δ*aroA*^, which was abrogated by asparagine or the asparaginase mutant (**Figure 7J**). This phenotype could also be replicated by the addition of the c-Myc inhibitor, which potently suppressed tumour stemness in a dose-dependent manner (**Figure 7K**). Finally, we validated these findings *in vivo*, and in keeping with the tumour organoid data and our previous study (Mackie et al., 2021), there was a significant reduction by STm^Δaro*A*^ in tumour stem cell markers by *Lgr5* and *Smoc2*, with a strong trend in *Vim*, a mesenchymal marker, which is a desirable outcome for tumour control (**Figure 7L**). Strikingly, oral treatment with STm^Δaro*A*/Δ*ansB*^ negated this intrinsic tumour control, as evidenced by an increase in expression of *Lgr5, Smoc2*, and *Vim*, to levels comparable to the PBS control group. In summary, T cells are metabolically inhibited by *Salmonella* due to asparaginase within the vector, however reversing the asparaginase activity comes at the cost of directly controlling of tumour stem cell growth and proliferation.

## Discussion

Bacterial cancer therapy occupies a unique niche in the cancer therapy landscape, being the historically oldest immunotherapy, yet at the forefront of biotechnological innovations. Bacterial vectors can be extensively engineered with additive functions (e.g. expression of cytokines (Yoon et al., 2017), checkpoint inhibitors (Pan et al., 2022), metabolic enzymes (Canale et al., 2021)), to great success in preclinical cancer models. To date, however, there is only one licensed BCT therapy in modern healthcare: the Bacillus Calmette–Guérin (BCG) vaccine, which is used to treat superficial bladder cancer (Pettenati and Ingersoll, 2018). Clinical trials using bacterial vectors—including *Salmonella—*have generally shown lacklustre results (Toso et al., 2002). It may be speculated that the naturally harboured immune evasion mechanisms these bacteria possess are a barrier to optimal therapeutic success, and one that is seldom explored.

Attenuated *Salmonella* is an ideal treatment vector for CRC for several reasons: (i) it matches the natural route of infection to the tumour location, and (ii) provides an opportunity for biological therapy in those patients whose cancer is poorly responsive to ICB – microsatellite instability^lo^, which comprises the majority of CRC patients (André et al., 2020; Zhao et al., 2019). The immune response to STm BCT has been thoroughly investigated, revealing the requirement of innate cells and innate immune adaptors for treatment efficacy (Chen et al., 2021; Johnson et al., 2021; Zheng et al., 2017). Yet while T cells have been shown redundant for treatment efficacy for two decades (Luo et al., 2001), canonical T cell activation and function is still assumed. This presumption is made despite the extensive immune evasion mechanisms harboured by *Salmonella* (Bernal-Bayard and Ramos-Morales, 2018; Wang et al., 2020).

Using a transgenic TCR reporter mouse that included temporal analysis of TCR signalling, we showed for the first time that T cells are suppressed during BCT, exhibiting defects in sustaining TCR signalling, cytokine production and proliferation. This was not due to problems in the TCR signalosome *per se*, but rather, a highly specific suppression of one master controller of T cell metabolism: c-Myc. Deletion of a single bacterial enzyme, or addition of a single amino acid, was sufficient to restore c-Myc expression and canonical T cell activation. In other words, T cells were poised to respond to TCR signalling, but prevented from sustaining an activation trajectory due to asparagine depletion. This effect is likely to be dependent on the local environment, in particular the presence of live STm.

This reinvigoration, however, came at the cost of intrinsic tumour control by STm. We have shown previously that STm can directly affect intestinal tumours by suppressing their stem cell-like characteristics, and also by imposing metabolic competition (Mackie et al., 2021). The data herein provide a mechanistic underpinning for these previous observations, and a target to improve therapeutic outcomes. c-Myc is a highly desirable oncotarget but one that is challenging for drug design due to the ubiquitous and essential nature of c-Myc in both tumour and normal cells (Madden et al., 2021). *Salmonella* as a tumour-homing vector, and one which can specifically invade *Lgr5^+^* cancer stem cells (Mackie et al., 2021), is uniquely placed to exploit this pathway in tumour control.

As to whether this ‘zero-sum game’ can be overcome—i.e. maintaining intrinsic antitumour effects while preserving T cell immunity—we theorise that this could be possible by combining STm BCT with transgenic CAR T-cell therapy, in which T cells are armoured against the effects of asparagine depletion by overexpression of asparagine synthetase or c-Myc itself. This will be the subject of further investigation by our laboratory.

One element that we did not resolve is the innate-like antiviral signature that T cells exhibited in the RNAseq analysis. However, these data are completely consistent with the later finding of diminished c-Myc protein by STm. Multiple components of the type I IFN pathway, such as IFN-β and IRF7, can antagonise c-Myc expression (Kim et al., 2016; Sarkar et al., 2006). This relationship is reciprocal, since c-Myc can in turn lead to degradation of key transcription factors in the type I IFN pathway, including STAT1 (Schlee et al., 2007). This early innate-like signature, therefore, is highly likely to reflect a rapid degradation of lymphocyte c-Myc by infected tumours.

We speculate that this study could have important implications for other bacterial vectors, namely the *Escherichia coli* strain Nissle (EcN). EcN is a probiotic bacterial strain that has been studied extensively in the prevention of CRC, and has recently been used in a study to modulate the metabolome of the TME (Canale et al., 2021). EcN contains a homologue of the *Salmonella* asparaginase that paralysed T cell metabolism in the present study – in fact, *E.coli* asparaginase is used as a chemotherapeutic. Countering this anti-T cell property—while sparing the direct antitumour effects—could unleash the full therapeutic potential of these bacterial vectors. In summary, we have shown for the first time that T cells are metabolically disabled in *Salmonella-based* BCT and proposed a model of asparagine depletion by the bacteria as the mechanism underpinning this. This asparagine depletion, however, is crucial for the direct anti-tumour effects of the bacteria. Anti-T cell and anti-tumour effects in BCT are therefore inexorably tied and must be disentangled for maximal treatment potential.

## Acknowledgements

We thank Professor Adam Cunningham (University of Birmingham, UK) for providing the auxotrophic aroA *Salmonella* strain, and Dr Rebecca Drummond (University of Birmingham, UK) and Dr Wei Li-Yu (University of Edinburgh, UK) for constructive feedback on the work. We also thank Professor Leslie Berg (University of Colorado, US) for providing the nuclei isolation protocol. Professor Anne Cooke (University of Cambridge, UK) generously provided functional antibodies. Work was funded by a CRUK Career establishment award (C61638/A27112, to K.M.M.), MRC CDA award (MR/V009052/1, to D.B.) and Biotechnology and Biological Sciences Research Council funding (BB/J013951/2 to M.O.).

## Materials and Methods

### Mice

Three strains of mice were used in this study: (i) C57BL/6 (Charles River, UK), (ii) *Nr4a3*-Timer (“Tocky”) (Bending et al., 2018a, 2018b) crossed with Great-Smart *ifng*-YFP reporter (Price et al., 2012), and (iii) *Nr4a1-Tempo* mice (Elliot et al., 2022). Males and females aged 6-32 weeks were used throughout the experiments, with age matching wherever possible. All animal experiments were approved by the Institutional Animal Care and Use Committees of RIKEN Yokohama Branch (計†2018-1(3)) and Yokohama City University (T-A-17-001) (Fig 7F) or the University of Birmingham local animal welfare and ethical review body and authorized under the authority of Home Office license P06118734 (held by K.M.M.) (all other experiments). Animal were maintained under SPF conditions with 12 hr day/night cycles and chow and water were fed ad libitum.

For use of *Nr4a3-Tocky* mice please contact Dr Masahiro Ono.

### Bacteria

*Salmonella enterica* serovar Typhimurium (SL3261) with a deletion in *aroA* (3-phosphoshikimate 1-carboxyvinyltransferase) was kindly provided by Professor Adam Cunningham (University of Birmingham, UK) (Flores-Langarica et al., 2015). The 4ansB mutant from STm 14028s single gene deletion library (Porwollik et al., 2014) was transduced into an *aroA* deleted STm UF20 strain (SR-11×3181 (Gulig and Doyle, 1993)) using P22 phage-mediated transduction (Schmieger, 1972). Both strains were grown in LB broth (Merck) overnight to saturation, followed by 1:20 sub-culture the following day until an OD_600_ of 0.7 was reached, indicating logarithmic growth. This sub-culture was then washed in PBS, concentrated, and used to inoculate either tumour organoids or mice. Colony-forming units (CFU) were enumerated in tumour lysates on LB agar (Merck) using standard microbiological techniques.

### Colitis-associated Colorectal Cancer (CAC) Model

C57BL/6 WT or *Nr4a3-Timer-GS* mice were used to induce CAC. WT mice were 7-8 week-old Mice (C57BL/6 WT or *Nr4a3*-Timer-GS) were injected with 10 mg/kg azoxymethane (Merck) via i.p. on day 0. On day 1, mice were given 3% dextran sodium sulfate (DSS) MW35-50,000 (APExBIO) in drinking water for ~7 days, followed by a two-week rest, and then two more cycles of 3% DSS. After a 1-2 week rest from DSS, mice were then orally gavaged with 5 x 10^9^ CFU bacteria in 100 μL volume, or PBS control, followed by a week rest, and then another dose of oral bacteria. A week later, mice were sacrificed, and tissue was collected for immunophenotyping. For tumour burden experiments, mice received 6 weekly doses of STm instead of 2, after which tumour burden was enumerated within the colon by inverted light microscopy.

Experiments using *Nr4a3*-Timer-GS were littermate mice and both males and females used. Experiments using purchased C57BL/6 mice used only female mice.

### Tumour Digestion and TIL Analysis

Tumours were dissected into tumour media (TM) containing ice-cold advanced DMEM-F12 (Gibco) containing 100 U/mL penicillin, 100 μg/mL streptomycin (Merck), 2% FCS (Gibco), 25 mM HEPES (Merck) and 50 μM β-mercaptoethanol (Merck). Tissue was then mechanically dissected into ~1 mm fragments using sterile scissors. Tumour fragments were then washed in 10 mL TM and the pelleted fragments were resuspended in pre-warmed 10 mL digestion buffer (DMEM-F12 containing 100 U/mL penicillin, 100 μg/mL streptomycin, 2% FCS, 0.1 mg/mL DNAse I (Roche) and 1 mg/mL collagenase D (Roche)) and digested in a 37°C orbital shaker under maximum rotation for 30-45 mins. Lastly, the digestion mixture was passed through a 70 μm cell strainer (BD Falcon), washed in FACS buffer, and processed for flow cytometry.

### Tumour Organoid Infection

Tumour organoids were derived from primary colonic tumours from the CAC model (C57BL/6 background) or small intestine and colonic tumours from adenomatous polyposis coli heterozygous mutant mice (Apc^Min/+^) (described in Mackie et al., 2021). Organoids were grown in 50 μL Matrigel (Corning) domes in 500 μL of tumour organoid media (DMEM-F12 containing 100 U/mL penicillin, 100 μg/mL streptomycin, 25 mM HEPES, 1:100 Glutamax (Gibco), 1:100 N-2 supplement (Gibco), 1:50 B27 supplement (Gibco), 1.25 mM N-acetylcysteine (Merck) and 50 pg/mL mEGF (Peprotech)). On day 3-4 of seeding, an overnight culture of STm was sub-cultured 1:20 in LB broth to an OD_600_ of approx.. 0.7. An inoculum of 5 μL containing ~1 x 10^8^ bacteria was added to each well, and then incubated at 37°C in 5% CO_2_ for 2 hours. Next, the media was aspirated and discarded, and each well was washed with fresh PBS twice. Finally, fresh tumour organoid media (500 μL) containing 50 μg/mL gentamicin (Gibco) was added, and the organoids were cultured for 24-48h. Tumour organoids were routinely passaged on a weekly basis and checked microscopically for organoid-like morphology. Heat-killed bacteria were prepared by heating the subculture to 95°C for 10 mins, and then adding 1 x 10^5^ bacteria (which is approximately equal to the CFU we recover from live infection) to the tumour media for the duration of the culture.

### Ex Vivo Tumour Culture

To culture primary tumours, tissue was excised from the CAC model as described and then cut into 1-2 mm pieces using sterile scissors. Equivalent amounts of tumour pieces were then resuspended in a 20-50 μL Matrigel dome and cultured in either tumour organoid media or TCM. Antibodies (1 μg/mL a-CD3 and 5 μg/mL a-CD28) were added to the appropriate wells in order to stimulate TILs, since these can penetrate the gel dome and access the tumour tissue (Voabil et al., 2021). For cytokine polyfunctionality assays, 10 μg/mL brefeldin A (Merck) was added during the 6h stimulation; for the *Nr4a3*-Timer experiment, antibodies alone were used for 16h. After stimulation, Matrigel was dissolved using 500 μL Cell Recovery Solution (Corning) for 30 mins at 4°C, followed by washing and then processing tumours as described for tumour digest and TIL analysis.

### Splenocyte Isolation

Splenocytes were isolated by mechanical disruption of the spleen, filtering through a 70 μm strainer, and then red blood cell lysis by ACK Lysis Buffer (Invitrogen) for 5 mins on ice. Cells were then re-filtered through a 70 μm strainer and kept on ice until further use.

### Tumour Organoid – Splenocyte/T-cell Co-culture

To assess T cell activation in the TME, splenocytes were directly cultured with tumour organoids after the 2h infection window with STm. Cells were seeded in the tumour wells at 1-2 x 10^6^ per well for the time indicated in the figure legends and the appropriate wells were stimulated with 1 μg/mL a-CD3 and 5 μg/mL a-CD28. Alternatively, cells were stimulated in 96-well round-bottom plates in the TCM from the end of a 48h tumour infection. This TCM was frozen at −20°C until use. Data points may indicate either individual mice or individual infections from pooled splenocytes, as specified.

To some conditions, the following were added at the indicated time-points: c-Myc inhibitor (10058-F4; Selleck Biochem), amino acids (10 mM glutamine, asparagine or aspartic acid – all from Merck), or D-glucose (50 mM; Merck).

### Proliferation Assay

To measure proliferation, splenocytes were stained with CellTrace Blue (Invitrogen) according to manufacturer instructions, and were then cultured in 96-well roundbottom plates in 100 μL TCM for 3 days ± activating antibodies (1 μg/mL a-CD3 and and 5 μg/mL a-CD28). Cells were then acquired on the flow cytometer and dye-low cells were indicative of proliferating cells.

### Enzyme-linked Immunosorbent Assay

Splenocytes (1 x 10^6^) were cultured in 96-well round-bottom plates in 100 μL TCM. Some conditions were activated using 1 μg/mL a-CD3 and 5 μg/mL a-CD28. Supernatants were collected after 24h and extracellular IL-2 was measured using the Biolegend ELISAMax Deluxe Kit (Biolegend), according to manufacturer instructions.

### General Flow Cytometry

Flow cytometry was performed in 96-well round-bottom plates. Cells were transferred to plates, and then washed in 100-200 μL FACS buffer (PBS containing 2% FCS, 2 mM EDTA (Merck) and centrifuged at 400 x *g* for 5 mins, after which the supernatant was discarded. Cells were then stained in the appropriate antibody master mix containing 1:1000 eFluor 780 Fixable Viability Dye (Invitrogen) and 1:500 Fc block (Prof. Anne Cooke, University of Cambridge). Cells were stained for 30 mins at 4°C while protected from light, then washed in FACS buffer and acquired on a BD Fortessa X20. UltraComp eBeads (Invitrogen) and unstained cells were used for compensation controls.

### Intracellular Flow Cytometry

For intracellular stains, cells were surface stained as described above, but then pelleted and resuspended in 100 μL fixative from the eBioscience Foxp3 / Transcription Factor Staining Buffer Set (Invitrogen) for 30 mins at 4°C while protected from light. Fixative was then washed off and discarded, and next cells were washed in FACS buffer followed by the kit permeabilisation buffer (Invitrogen). Cells were then resuspended in a staining mixture made-up in permeabilisation buffer and stained for 30-60 mins at 4°C in complete darkness. Next, cells were washed in permeabilisation buffer, and then in FACS buffer before data acquisition. However, antibodies requiring a secondary conjugation were instead stained for another 30 mins in permeabilisation buffer as an extra step, before washing and acquisition.

Flow cytometry of c-Myc in tumour organoids was performed by first dissolving Matrigel using Cell Recovery Solution (Corning), and then digesting the organoids into single cells using 1mL TryplE per condition at 37C for 5 minutes, followed by washing and proceeding as normal. All flow cytometry steps before fixation with tumour organoids were performed in the presence of ROCK inhibitor (Y-27632) used at 5 μM.

### PhosFlow

To stain phosphorylated epitopes, cells were stained according to surface stain procedures, but transferred to 5 mL polystyrene tubes and then washed twice in excess PBS. Cells were then resuspended in 1:1 PBS-diluted Fixation Buffer (Biolegend), comprising ~2% PFA, and incubated at 37°C in a water bath with gentle agitation. Following mild fixation, cells were centrifuged (1000 x *g* for 5 minutes), PFA was discarded, and then cells were washed twice with PBS containing 2% FCS. The cell pellet containing residual buffer was vortexed and then 1 mL TruePhos Fixation Buffer (Biolegend) was added drop-by-drop while under constant vortexing. Samples were then stored at −20°C in the dark for 1h. Samples were next centrifuged and the fixation buffer discarded, followed by two washes in PBS/FCS. Finally, cells were stained with phospho-antibodies in PBS/FCS for 30-45 mins at 4°C in complete darkness, followed by a wash and then acquisition on flow cytometer.

### Calcium Flux Assay

Splenocytes (1 x 10^6^) were cultured with TCM from infected or non-infected tumours for 24h. Next, cells were surface stained for flow cytometry in 5 mL polystyrene tubes as described, washed in PBS, and then stained with 2.5 μM Fura-Red and Flura-4 mixture (both Invitrogen) for 30 mins at 37°C. Cells were then washed twice in PBS containing 2% FCS to quench any extracellular dye, rested for 10 minutes, followed by one wash in pure PBS. Cells were then acquired on the flow cytometer for ~25 seconds before the addition of 1 μg/mL ionomycin (Merck), and calcium flux was measured for ~1.5 mins.

### Nuclear Flow Cytometry

Analysis of transcription factors within T cell nuclei was performed according to an established protocol (Gallagher et al., 2021, 2018). Bulk T cells were first negatively isolated using either MojoSort (Biolegend) or MACS separation (Miltenyi Biotech) to high purity (> 97%), according to manufacturer instructions. Next, T cells were stimulated in TCM from non-infected or STm-infected tumours for 24h. One hour before the end of culture, 1 μg/mL a-CD3 and 5 μg/mL a-CD28 were added to stimulated conditions. After stimulation, cells were centrifuged, media was discarded, and the plate was put on ice. Sucrose buffer (200 μL/well) containing 10 mM HEPES, 8 mM MgCl_2_ (Merck), 320 mM sucrose, 1 x EDTA-free protease inhibitor (Roche) and 0.1% Triton-X100 (Merck) was added to each well, and cells were incubated for 15 mins. The plate was then centrifuged at 2000 *x* g for 5 minutes while chilled and the supernatant was discarded. Next, cells were washed twice in the sucrose buffer (lacking Triton-X100), followed by fixation in 4% PFA diluted in sucrose buffer (again lacking Triton-X100) for 30 mins on ice, protected from light. Cells were washed and supernatant discarded. Cells were then washed again in FACS buffer containing 8 mM MgCl_2_, followed by a wash in permeabilisation buffer (FACS buffer with 0.3% Triton-X100 and 8 mM MgCl_2_). Cells were stained in antibodies diluted in permeabilisation buffer for 1h, before washing with magnesium-supplemented FACS buffer and acquisition on the flow cytometer. Cells were stained for β-tubulin to check purity of nuclei, which also served as a gate to exclude any cytoskeletal debris.

### RNA extraction and RNAseq

Splenocytes were activated in TCM from infected or non-infected tumours as described, and then at 4h or 16h CD4 T cells were fluorescently sorted to high purity on a FACSAria. RNA was extracted from lysates using the Arcturus Picopure RNA kit (ThermoFisher) according to the manufacturer’s instructions. 5 ng of RNA was used for generation of sequencing libraries using the Quantseq 3’ mRNA-seq Library Preparation kit FWD (Lexogen). Unique Molecular Identifiers (UMIs) were used for the evaluation of input and PCR duplicates and to eliminate amplification bias. Libraries were normalised and pooled at a concentration of 4 nM for sequencing. Libraries were sequenced using the NextSeq 500 using a Mid 150v2.5 flow cell. Cluster generation and sequencing was then performed and FASTQ files generated. FASTQ files were then downloaded from the Illumina base space and uploaded to the BlueBee cloud for further analysis (Lexogen). FASTQ files were merged from the 4 lanes to generate final FASTQ files which were loaded into the BlueBee QuantSeq FWD pipeline and aligned to the GRCm38 (mm10) genome. HTSeq-count v0.6.0 was used to generate read counts for mRNA species and mapping statistics. Raw read counts in the .txt format were used for further analysis using DESeq2 (Love et al., 2014) in R version 4.0. A DESeq dataset was created from a matrix of raw read count data. Data were filtered to remove genes with fewer than 10 reads across all samples. Log2 fold change estimates were generated using the DESeq algorithm and shrinkage using the ashr algorithm (Stephens, 2017) to estimate log2 fold changes (lfc). Differentially expressed genes (DEGs) were selected based on an adjusted *p* value of < 0.05, and then gene lists filtered to identify genes with an estimated lfc greater or less than 1.5. Normalised read counts were transformed using the regularised log (rlog) transformation. Heatmap analysis was performed on the rlog transformed data using the R package gplots. For KEGG pathway analysis clusterProfiler (Yu et al., 2012), DOSE (Yu et al., 2015), and biomaRt (Durinck et al., 2009) packages were used.

### ECAR and OCR analysis

To measure extracellular acidification (ECAR) and oxygen consumption rate (OCR), splenocytes (2 x 10^6^/well) were activated in technical triplicate from each mouse for 48h in TCM from non-treated or STm-infected tumour organoids. Cells were then washed and resuspended in “Seahorse XF” serum-free RPMI (Agilent), seeded onto pre-coated (poly-d-lysine; Invitrogen) Seahorse plates and analysed on a Seahorse XFe96 metabolic extracellular flux analyser as previously described (Gudgeon et al., 2022). Glucose (10mM Sigma) was first added and then Oligomycin (1 μM; Merck), BAM-15 (3 μM; Merck), rotenone (2 μM; Merck) and antimycin A (2 μM; Merck) were injected to disrupt various elements of the metabolic pathways. Basal OCR was calculated as the mean of the initial 3 measurements minus the mean of the 3 measurements after rotenone/antimycin A injection. ATP-coupled OCR was calculated as the basal OCR minus the mean of the 3 measurements after oligomycin injection. Maximal OCR was calculated as the mean of the 3 measurements after BAM-15 injection minus the mean of the 3 measurements after rotenone/antimycin A injection. Spare respiratory capacity was calculated as maximal OCR minus basal OCR. Glycolysis was calculated as the mean of the 3 measurements after glucose injection minus the mean of the initial 3 measurements. Glycolytic capacity was calculated as the mean of the 3 measurements after oligomycin injection minus the mean of the initial 3 measurements. Glycolytic reserve was calculated as the glycolytic capacity minus glycolysis.

### Tumour Organoid qPCR

Tumour organoids were infected as previously described and dissociated using Cell Recovery Solution and TryplE. qPCR was performed as previously described (Mackie et al., 2021). Briefly, RNA was isolated from single cell suspensions using a commercial kit (Qiagen RNeasy Mini Kit, Qiagen) according to manufacturer instructions. m-MLV and oligoDTs (Merck) were used to generate cDNA, and qPCR was performed using a Roche LightCycler 480 with SYBR green system (Takara). Below are the primers used for this study:

*Lgr5:* 5’-CGGAAAGTGGAATCCTTGCA, 3’-CACATCGATCTGGACATGCTGT
*Vim:* 5’-CGGAAAGTGGAATCCTTGCA, 3’-CACATCGATCTGGACATGCTGT
*Smoc2:* 5’-CCCTCAGAAGCCACTCTGTG, 3’-ACTTGCTGGAACTCCTTCCG
*Gapdh:* 5’-TGTGTCCGTCGTGGATCTGA, 3’-TTGCTGTTGAAGTCGCAGGAG

### MTT Assay

Tumour organoids were infected as previously described and dissociated using Cell Recovery Solution and TryplE, and then seeded into 96-well flat bottom plates at 4 x 10^4^/well in 10 μL Matrigel domes. 150 μL tumour organoid media was placed on top of each dome. After 24h, 50 μL media was removed and replaced with 50 μL MTT reagent (Abcam) which was incubated for 1-3 hours at 37°C and 5% CO_2_. Upon the appearance of purple crystals, the assay was halted by addition of 4 mM HCl, 0.1% NP40 in isopropanol. Optical density was then measured by plate-reader at OD560. Results were performed in triplicate and averaged.

### Antibodies

The following antibodies were used in this study: β-tubulin-AF488 (9F3; CellSignaling), CD3ε (145-2C11; Biolegend), CD28 (37.51 from hybridoma; Prof Anne Cooke, University of Cambridge), CD4-BUV395 (GK1.5; BD), CD4-BUV737 (GK1.5; BD), CD4-BV421 (GK1.5; Biolegend), CD8a-PE/Cy7 (53-6.7; Biolegend), CD8α-AF700 (53-6.7; Biolegend), CD45-BUV395 (30-F11; BD), CD45-FITC (30-F11; Biolegend), CD45-PerCP/Cy5.5 (30-F11; Biolegend), CD69-AF700 (H1.2F3; Biolegend), CD44-PE/Cy7 (IM7, Biolegend), CD62L-APC (MEL-14; Biolegend), TCRβ-PerCP/Cy5.5 (H57-597; Biolegend), TCRβ-AF700 (H57-597; Biolegend), PD-1-PE/Cy7 (29F.1A12; Biolegend), CD90.2-APC (53-2.1; Biolegend), Glut1 (SA0377; Invitrogen), Glut3 (JA50-31; Invitrogen), p-Erk1/2-T202/Y204-AF647 (4B11B69; Biolegend), NFAT1-AF647 (D43B1; CellSignaling), NF-κB-PE (D14E12; CellSignaling), p-Akt-S473-PE (SDRNR; eBioscience), p-mTOR-S2448-PE/Cy7 (MRRBY; eBioscience), p-p70-S6K-T421/S424 (polyclonal rabbit; CellSignaling), IL-2 (JES6-5H4; Biolegend), TNFa-BV605 (MP6-XT22; Biolegend), IFN-γ-AF647 (XMG1.2; Biolegend), c-Myc (E5Q6W; CellSignaling), p-Lck-Y394 (A18002D; Biolegend) and F(ab’)2-Goat anti-Rabbit IgG (H+L) Cross-Adsorbed-APC (Invitrogen). eFluor-780 Fixable Viability Dye (Invitrogen) was used for discriminating live/dead cells.

### Software and Statistical Analysis

Data were analysed primarily using FlowJo version 9, GraphPad Prism version 9 and Microsoft Excel. RNAseq analysis was performed using software described in the relevant section. Graphical figures were made using BioRender (BioRender.com) and Microsoft Powerpoint. Statistical analyses are specified in the relevant figure legends, with appropriate post-tests to control for multiple testing. Data from individual mice in some figures were excluded if recovered tumour cell populations were of insufficient size. Statistical significance was determined as being *p* ≤ 0.05.

## Supplementary Figures

**Supplementary Figure 1.**
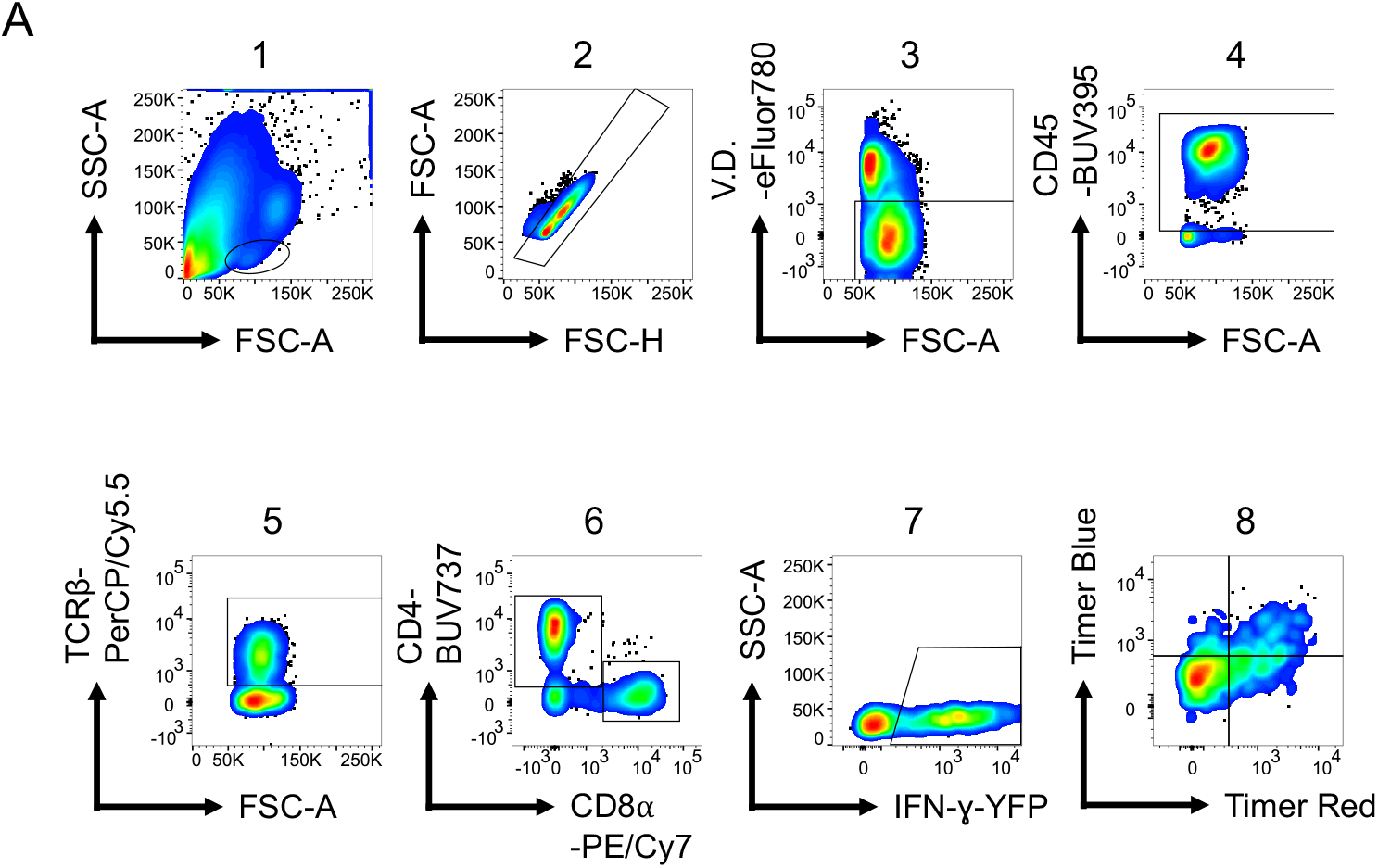
A representative gating strategy for analysis of TILs from the CAC model. Cells were gated by size to exclude epithelial debris (1) → single cells (2) → live cells (3) → leukocyte lineage (4) → T cells (5) → CD4 or CD8 (6) → IFN-g positive (7) → *Nr4a3*-Timer expression.

**Supplementary Figure 2.**
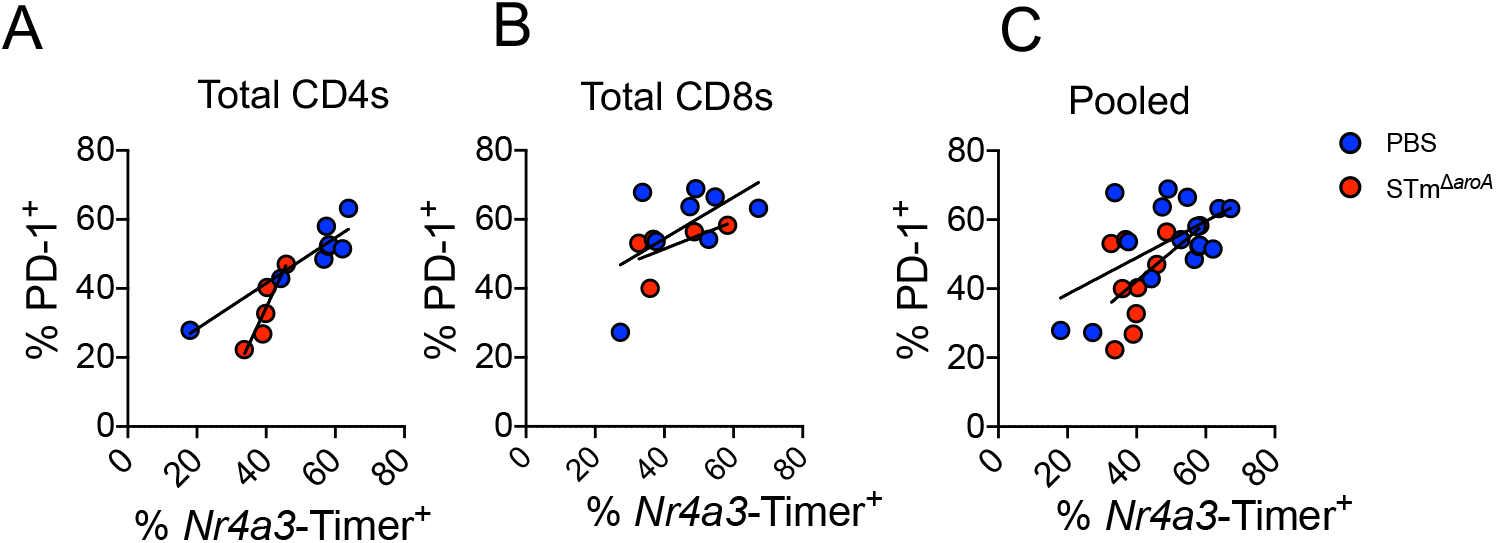
Correlation between PD-1 expression and *Nr4a3-Timer* positivity. TILs were analysed from digested primary tumours and total CD4s (A), CD8s (B) or pooled T cells (C) were analysed for correlation between PD-1 positivity and *Nr4a3-Timer* positivity. Linear regression was applied to both data sets; line depicts best fit of the regression. Each data point represents a single mouse.

**Supplementary Figure 3.**
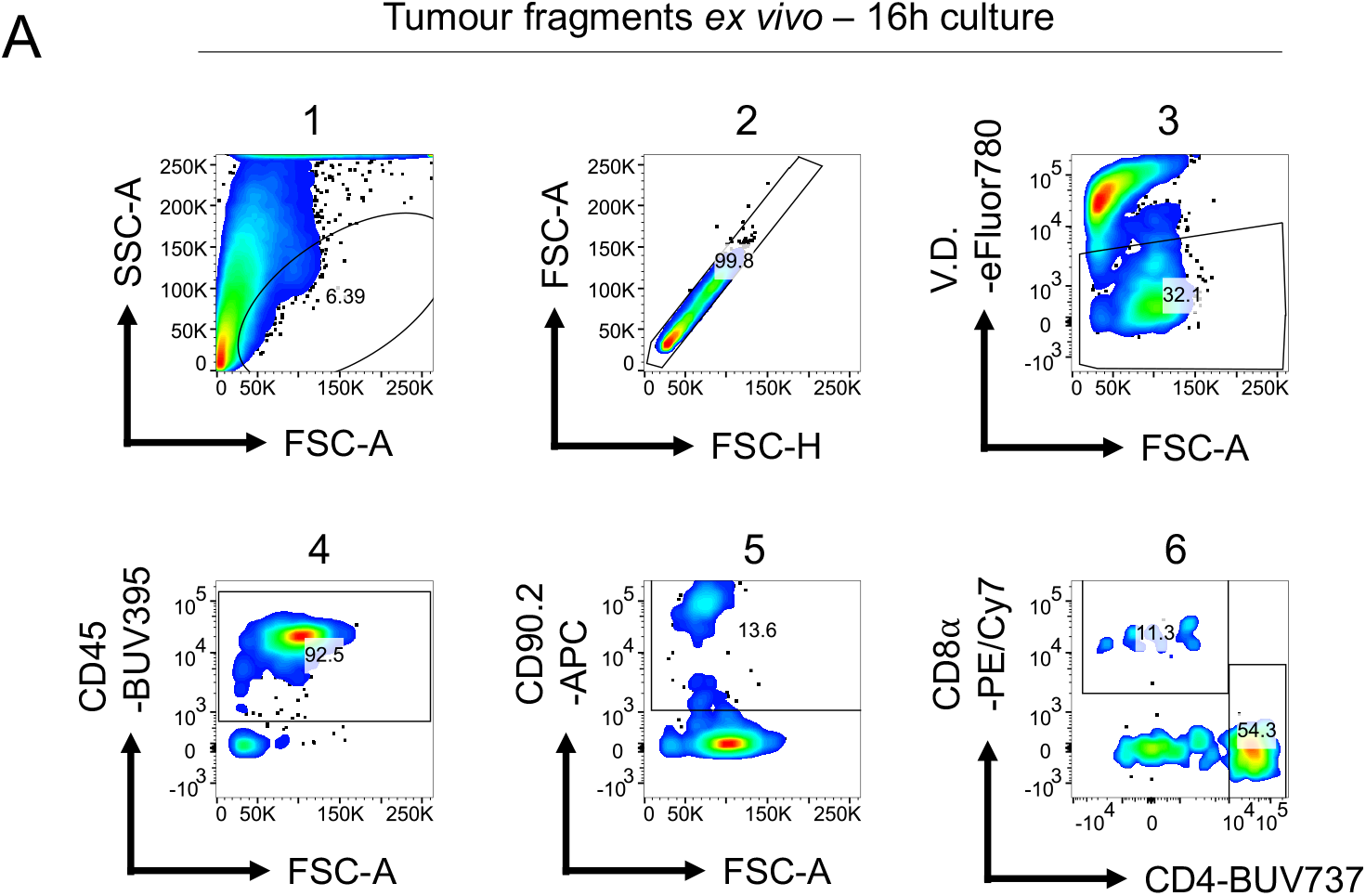
(A) Representative gating strategy depicting analysis of TILs that had been cultured *ex vivo* in Matrigel for 16h after dissection. Cells were stimulated with a-CD3/a-CD28 antibodies (1 μg/mL and 5 μg/mL, respectively) for 16h and then processed for flow cytometry. Cells were gated by size to exclude epithelial debris (1) → single cells (2) → live cells (3) → leukocyte lineage (4) → T cells (5) → CD4 or CD8.

**Supplementary Figure 4.**
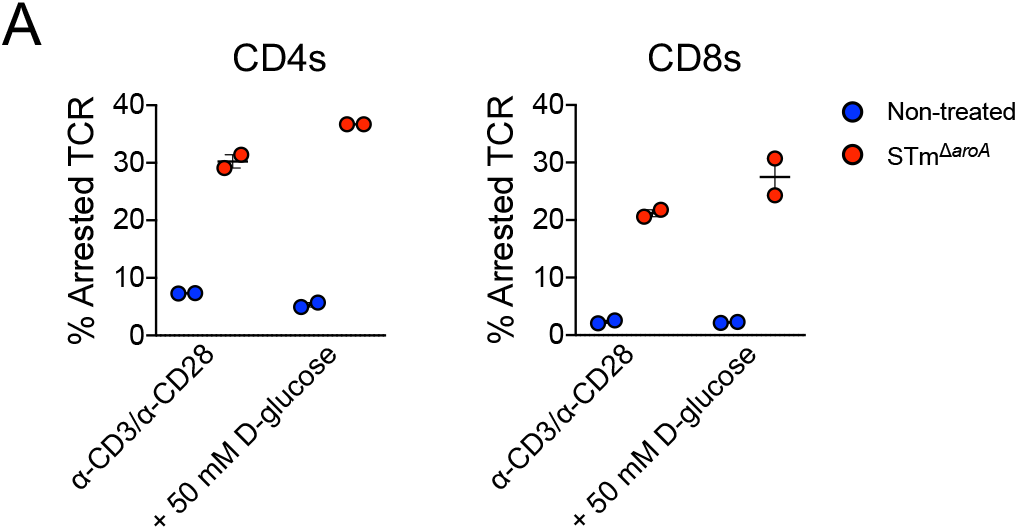
(A) Splenocytes were co-cultured with infected or non-infected tumours for 24h in the presence of a-CD3/a-CD28 antibodies (1 μg/mL and 5 μg/mL, respectively). During the final 6h, 50 mM glucose was spiked into the culture in an attempt to rescue arrested TCR activation, as measured by *Nr4a3*-Timer Arrested TCR signal, i.e. *Nr4a3*-Timer Blue^neg^Timer Red^pos^. Cells were then aspirated and tested for activation by flow cytometry. Data show two independent infections; each data point represents splenocytes tested with an independent tumour infection.

**Supplementary Figure 5.**
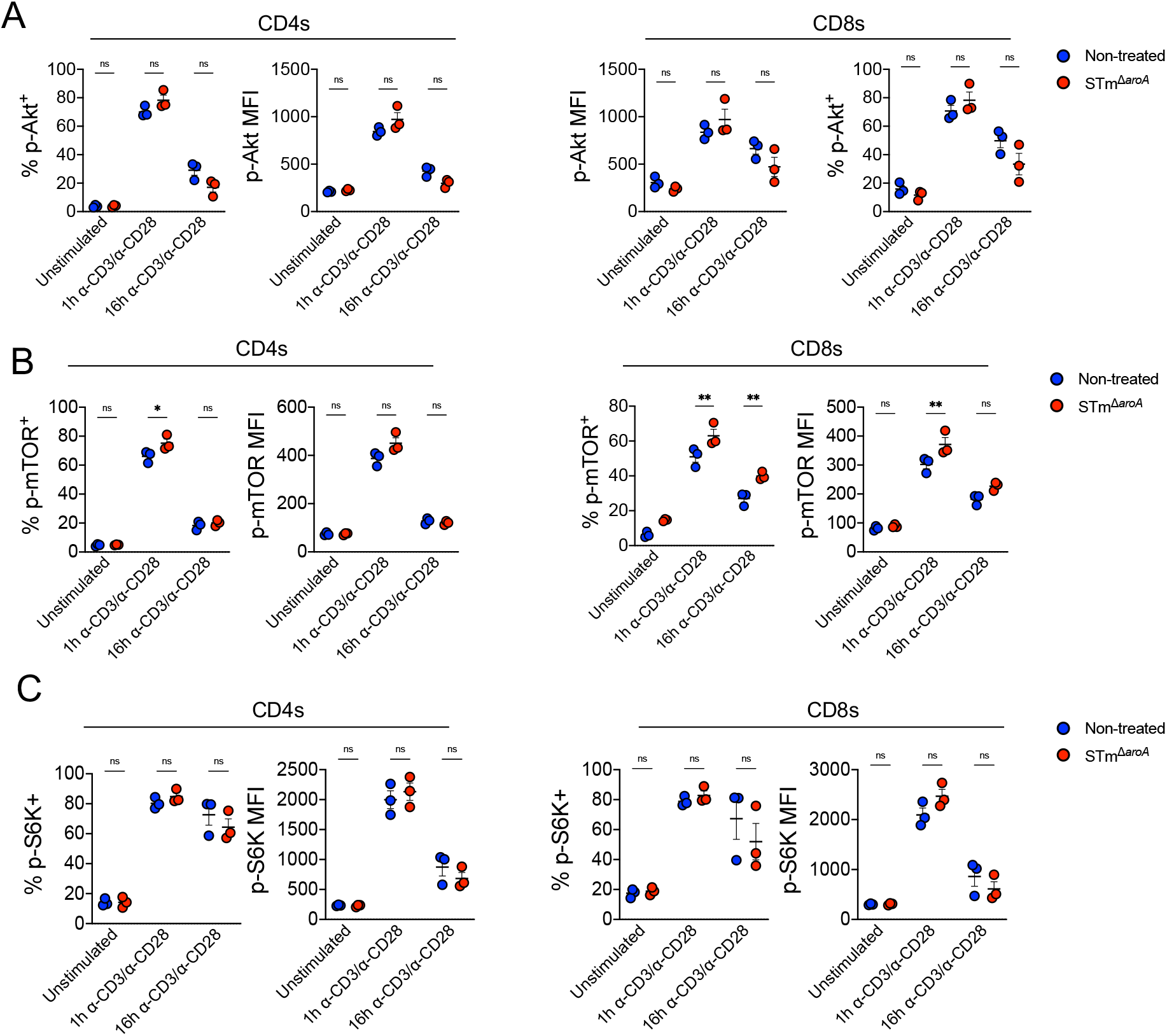
Detailed analysis of PhosFlow signalling pathways as shown by representative plots in Figure 6E. Cells were stimulated in NT and STm TCM for 1h or 16h and processed for PhosFlow as previously described, showing p-Akt-S473 (*A*), p-mTOR-S2448 (*B*) and p-70-S6K-T421/S424 (*C*). Bars depict means ± SEM. Statistical significance was tested by two-way ANOVA with Sidak’s post-test (non-treated vs STm). Data are derived from *n* = 3 mice testing pooled TCM from two tumour infections.* = *p* < 0.05, ** = *p* < 0.01, *** = *p* < 0.001, **** = *p* < 0.0001.

**Supplementary Figure 6.**
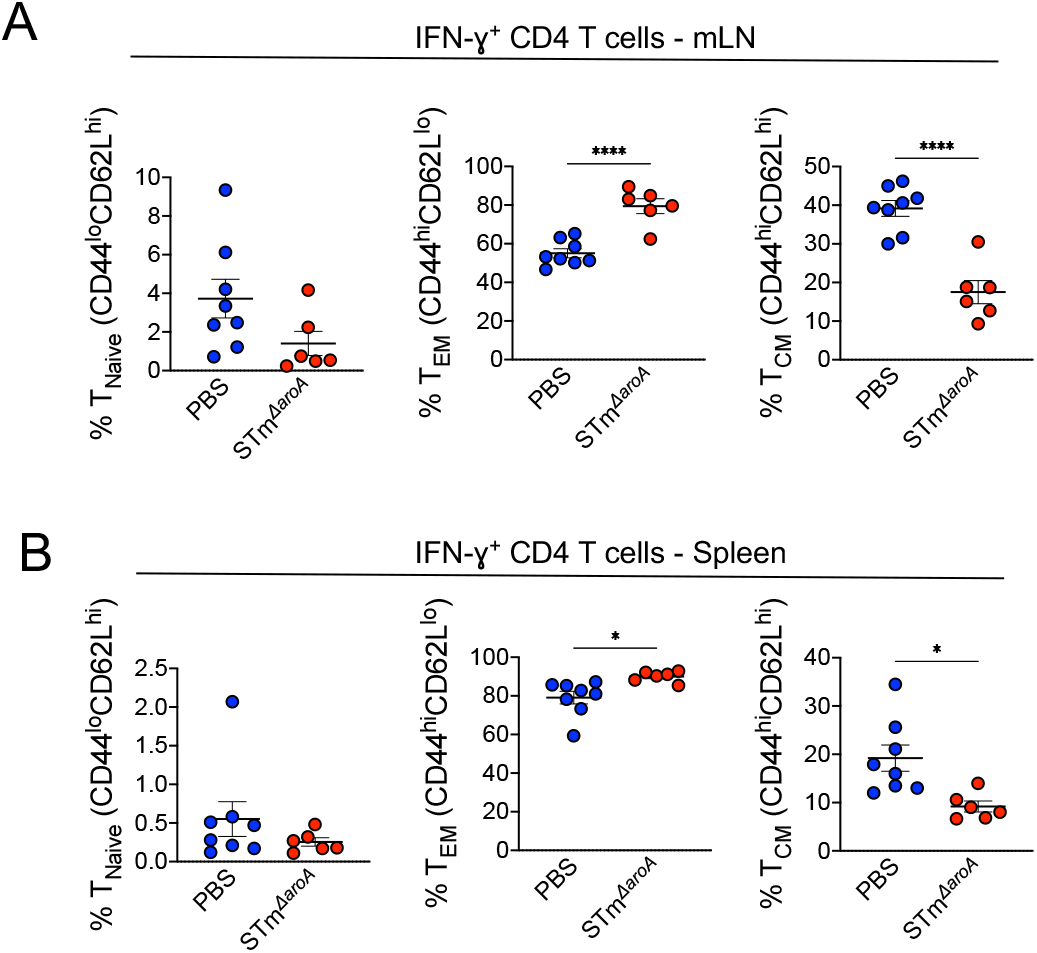
Tumours were induced in mice using the CAC model, followed by two rounds of oral STm treatment as previously outlined. One week after the final dose, mice were culled and mLN and spleens were extracted, followed by preparation of single cell suspensions and FACS staining for T cell memory subsets within the IFN-γ^+^ CD4 (*A*) and CD8 T cell (*B*) populations. Cells were phenotyped based upon expression of CD44 and CD62L. Bars depict means ± SEM. Each data point represents one mouse. Statistical significance was tested by unpaired two-tail t-test.. * = *p* < 0.05, ** = *p* < 0.01, *** = *p* < 0.001, **** = *p* < 0.0001.

**Supplementary Figure 7.**
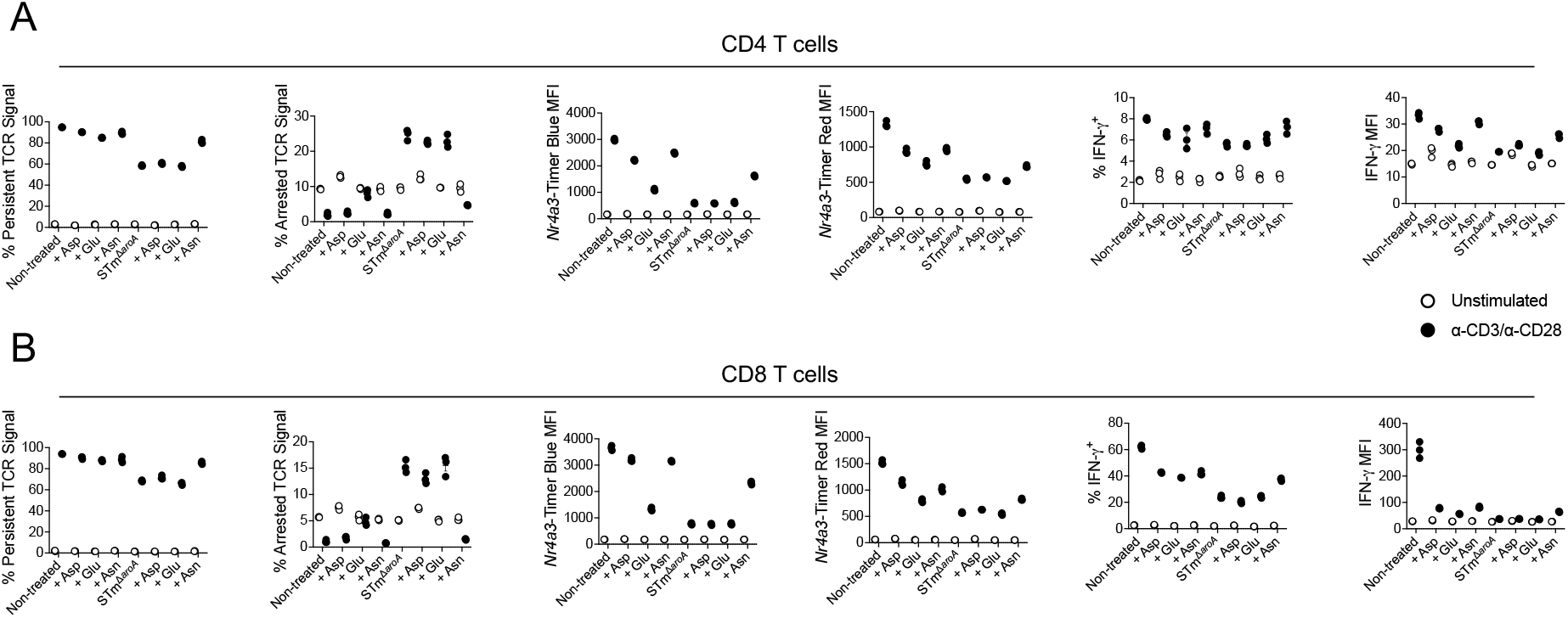
Splenocytes were cultured in TCM from either non-infected or infected-tumours and activated for 24h with a-CD3/a-CD28 antibodies (1 μg/mL and 5 μg/mL, respectively). To some cultures, Asp/Glu/Asn were added at 10 mM at the beginning of the culture, or else a vehicle control (H20) was used. Various metrics of either *Nr4a3-Timer*, indicative of TCR signalling, or IFN-g expression were quantified by flow cytometry. Data are from *n* = 3 mice, using pooled TCM from two tumour infections. Bars depict means ± SEM.

